# SIX2 and SIX3 coordinately regulate functional maturity and fate of human pancreatic β cells

**DOI:** 10.1101/2020.12.03.411033

**Authors:** Romina J. Bevacqua, Jonathan Y. Lam, Heshan Peiris, Robert L. Whitener, Seokho Kim, Xueying Gu, Mollie S.H. Friedlander, Seung K. Kim

## Abstract

The physiological functions of many vital tissues and organs continue to mature after birth, but the genetic mechanisms governing this postnatal maturation remain an unsolved mystery. Human pancreatic β-cells produce and secrete insulin in response to physiological cues like glucose, and these hallmark functions improve in the years after birth. This coincides with expression of the transcription factors SIX2 and SIX3, whose functions in native human β-cells remain unknown. Here, we show that shRNA-mediated *SIX2* or *SIX3* suppression in human pancreatic adult islets impairs insulin secretion. However, transcriptome studies revealed that *SIX2* and *SIX3* regulate distinct targets. Loss of *SIX2* markedly impaired expression of genes governing β-cell insulin processing and output, glucose sensing, and electrophysiology, while *SIX3* loss led to inappropriate expression of genes normally expressed in fetal β-cells, adult a-cells and other non-β-cells. Chromatin accessibility studies identified genes directly regulated by SIX2. Moreover, β-cells from diabetic humans with impaired insulin secretion also had reduced *SIX2* transcript levels. Revealing how *SIX2* and *SIX3* govern functional maturation and maintain developmental fate in native human β-cells should advance β-cell replacement and other therapeutic strategies for diabetes.

## Introduction

Development of vital organs, like the brain and pancreas, includes a prenatal stage when the embryonic organ anlage, specification and expansion of major differentiated cell types, organ morphology and anatomic position are established, followed by a postnatal stage when differentiated cell types acquire mature physiological functions and refine their cellular interactions (Reinert et al. 2014; Kim et al. 2020). Improved understanding of the mechanisms underlying postnatal “functional maturation” in cells like pancreatic islet cells could accelerate efforts to improve therapies for diabetes, including production of replacement human β-cells from renewable sources (Bakken et al. 2016; Arda et al. 2016; Arda et al. 2018; Sneddon et al. 2018).

During prenatal and neonatal stages, islet β-cells transiently proliferate and expand. In childhood and thereafter, β-cells reduce proliferation (Teta et al. 2005; Meier et al. 2008; Wang et al. 2016a), increase insulin production and enhance glucose-dependent insulin secretion, all hallmarks of mature β-cell function (Arda et al. 2016; Aguayo-Mazzucato et al. 2011; Artner et al. 2007; Artner et al. 2010). The genetic and molecular mechanisms governing this age-dependent β-cell functional maturation are intensely sought. Prior studies in mice suggest that transcription factors (TFs) regulate functional maturation of β-cells (Lantz et al. 2004; Schaffer et al. 2013; Yoshihara et al 2016; Aguayo-Mazzucato et al. 2018). However, less is known about the role of TFs in human β-cell maturation, reflecting the challenges of studying postnatal development in native human β-cells. Mutations in genes encoding TFs including *PDX1, NEUROD1, MAFA, GATA6* and *RFX6* have been linked to monogenic forms of diabetes mellitus and impaired β-cell function in humans, suggesting roles in β-cell maturation (reviewed in Urakami et al. 2019; Sellick et al. 2004; Patel et al. 2017; Allen et al. 2011; Iacovazzo et al 2018). Thus, TFs may represent a class of factors governing age-dependent postnatal β-cell functional maturation in humans.

We and others have found that SIX2 and SIX3, members of the *<u>si</u>ne oculus* homeobo<u>x</u> family of TFs, are first expressed in the β-cells of children by 9-10 years of age (Arda et al. 2016; Blodgett et al. 2015; Cyphert et al. 2019; Wang et al. 2016b), followed by increased expression in adulthood. Moreover, neither *SIX2* nor *SIX3* mRNA are detectable in human a-cells (Arda et al. 2016; Blodgett et al. 2015; Wang et al. 2016b). *SIX2* and *SIX3* are encoded by linked genes on human chromosome 2 (OMIM: 604994 and 603714). While they show high homology in their N-terminal SIX domain and their central homeodomain (HD), other portions of SIX2 and SIX3 differ (Kawakami et al. 2000), and could mediate distinct interactions. For example, SIX3, but not SIX2, interacts with members of the *groucho* family of corepressors (Kobayashi et al. 2001; López-Ríos et al. 2003). In addition, SIX2 has roles in development of kidneys, skull, stomach and other organs (Kobayashi et al. 2008; He et al. 2010; Self et al. 2009; Park et al. 2012), while SIX3 has roles in forebrain and eye development (Jeong et al. 2008; Liu et al. 2010).

*SIX2* and *SIX3* have largely non-overlapping tissue expression patterns that correspond with their distinct roles in human organogenesis. In the pancreas, however, both *SIX2* and *SIX3* are expressed in human β-cells, with coincident onset of postnatal expression and a shared *cis*-regulatory element that coregulates islet *SIX2* and *SIX3* expression (Spracklen et al. 2018). This element encompasses single nucleotide polymorphisms (SNPs) associated with type 2 diabetes (T2D) and fasting hyperglycemia (Kim et al. 2011; Spracklen et al. 2020). Nevertheless, it remains unknown whether SIX2 and SIX3 are required for normal adult β-cell function, what β-cell genes these TFs regulate, and whether their expression is altered in diabetes.

Prior studies suggest that *SIX2* or *SIX3* might regulate genes with roles in restricting native β-cell proliferation, and enhancing insulin production and secretion. The onset of *SIX2* and *SIX3* expression in postnatal human β-cells temporally coincides with age-dependent enhancement of insulin production and secretion (Arda et al. 2016; Blodgett et al. 2015). Moreover, mis-expression of *SIX3* in immature islets from children stimulated glucose-dependent insulin secretion (Arda et al. 2016). Neither *SIX2* nor *SIX3* is expressed in mouse islets (Benner et al. 2014; Baron et al. 2016); thus, loss-of-function studies that clearly identify requirements for SIX2 or SIX3 in β-cell function will require studies in native human islets, or alternative experimental systems. Attempts to generate insulin-secreting β-cells from human stem cells have only produced immature progeny that express *SIX2,* often at low levels, and fail to express *SIX3* (Sneddon et al. 2018; Nair et al. 2019; Veres et al. 2019; Velazco-Cruz et al. 2020). To overcome these limitations, here we investigated *SIX2* and *SIX3* in adult human islets, using recently-described genetic methods permitting targeted loss-of-function in primary human islets (Peiris et al. 2018). Controlled islet cell dispersion and re-aggregation to develop human ‘pseudoislets’ (Arda et al. 2016; Scharp et al. 1980; Tze et al. 1982) allowed us to achieve shRNA-mediated *SIX2* or *SIX3* suppression in human adult β-cells. Collectively, these studies demonstrate that *SIX2* and *SIX3* coordinately regulate distinct genetic programs in human β-cells, including expression of target genes governing mature β-cell function and maintaining β-cell fate. Moreover, we show that *SIX2* expression is reduced in β-cells purified from human donors with T2D and impaired islet insulin secretion, suggesting roles for *SIX2* in the pathogenesis of β-cell defects in T2D.

## Results

### Reduced SIX2 or SIX3 expression impairs regulated insulin secretion by human islets

We used RNAi-based strategies to reduce *SIX2* or *SIX3* mRNA levels in primary human adult islets. After dispersion of primary islets, lentiviral transduction was used to achieve shRNA-mediated suppression of *SIX2* or *SIX3* (hereafter, *SIX2*^kd^ or *SIX3*^kd^) and to simultaneously express a *GFP* transgene (Peiris et al. 2018), followed by reaggregation into pseudoislets (Fig. 1A-B; Methods). We used immunostaining to detect β-cells (Insulin, INS), a-cells (Glucagon, GCG) and δ-cells (Somatostatin, SST) in pseudoislets, and verified re-aggregation of these principal islet cell-types in appropriate proportions (Supplemental Fig. 1). Five days after lentiviral infection, we confirmed significant reduction of *SIX2* or *SIX3* in pseudoislets by qRT-PCR (Fig. 1C-D; Methods). By contrast, we did not detect altered *SIX3* mRNA levels after *SIX2*^kd^ or altered *SIX2* mRNA levels after *SIX3*^kd^ (Fig. 1E-F), providing evidence that *SIX2* and *SIX3* expression are not mutually cross-regulated.

**Figure 1.**
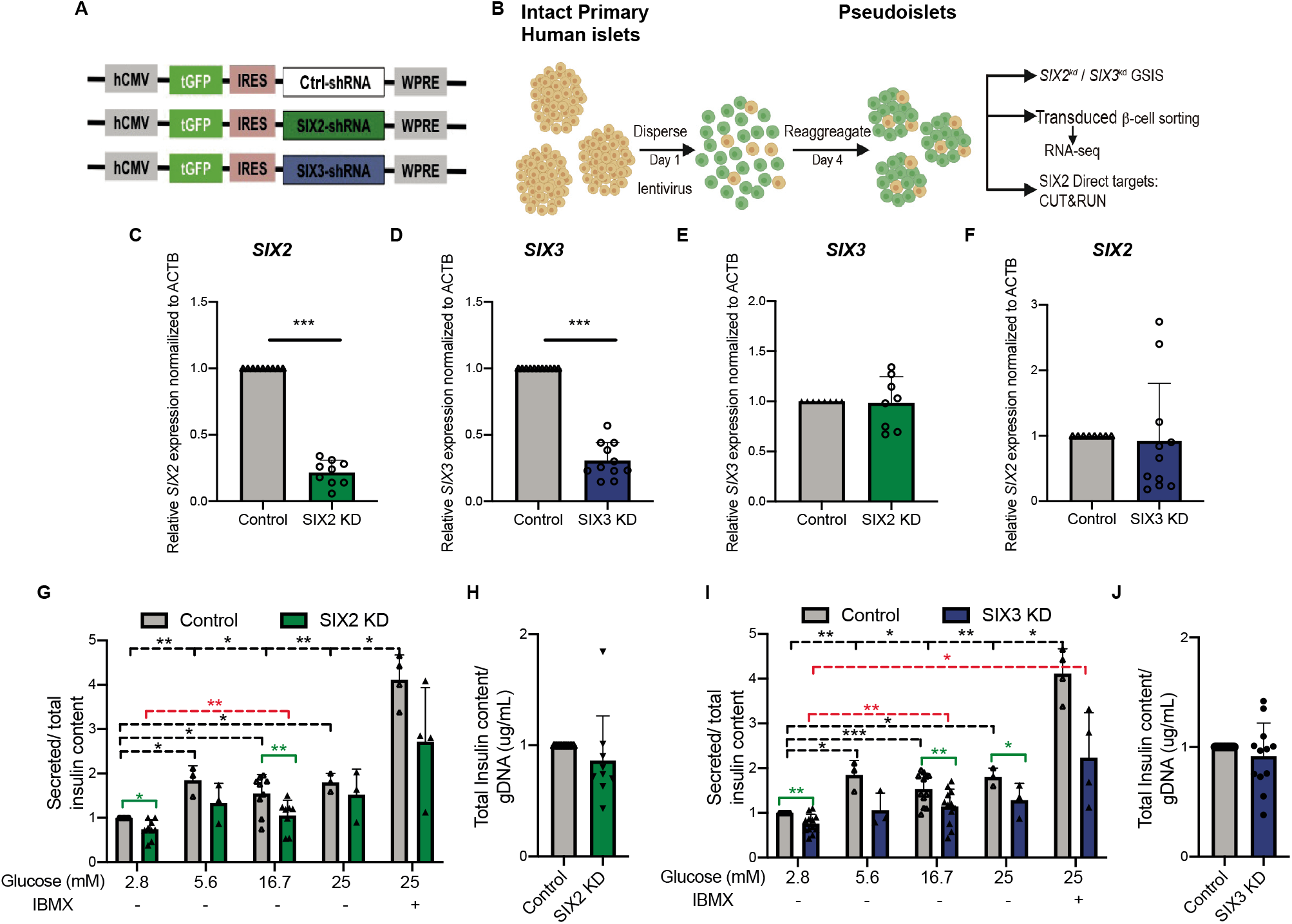
shRNA-mediated suppression of *SIX2* and *SIX3* in primary human islets results in impaired glucose stimulated insulin secretion. (A) Schematics of the lentiviral constructs coding for a short hairpin RNA (shRNA) and GFP. (B) Schematic detailing the pseudoislet technique. (C) *SIX2* mRNA expression in primary human islets control (grey bar) or *SIX2^kd^* (green bar) (n=9 independent donor repetitions). (D) *SIX3* mRNA expression in primary human islets control (grey bar) or *SIX3^kd^* (blue bar) (n=11 independent donor repetitions). (E) *SIX3* mRNA were not affected by *SIX2^kd^* (n=8 independent donor repetitions) and (F) *SIX2* mRNA levels were not affectsd by *SIX3*^kd^ (n=11 independ tint d onor repetitions). (G,I) *In vitro* glucose-stimulated insulin secretion from human pseudoislets control, (G) *SIX2^kd^* (n=9 independent donor repetitions) or (l) *SIX3^kd^* (n=12 independent donor repetitions). Secreted insulin normalized to insulin content, black lines highlight significant differences within the control; red, within the KD groups, and green between control and KD conditions. (H, J) Total insulin from human pseudoislets after transduction with (H) *SIX2^kd^* (n=9 independent donor repetitions) or (J) *SIX3^kd^* (n=12 independent donor repetitions). Data presented as mean, error bars represent the standard error. Two-tailed tests sts used to generate *P* values. * *P*< 0.05, ** *P*< 0.01 and *** *P*<0.0001.

We subsequently assessed glucose-stimulated insulin secretion (GSIS) following *SIX2^kd^* or *SIX3*^kd^ in primary human pseudoislets. Like in our prior studies (Peiris et al 2018), control pseudoislets infected with lentivirus expressing non-targeting shRNA (‘Control’) showed a significant increase in insulin secretion after a glucose step increase from 2.8 mM to 5.6 mM, 16.7 mM, 25 mM or 25 mM glucose supplemented with the secretion potentiator IBMX (Fig. 1G,I). By comparison, insulin secretion by pseudoislets after S*IX2*^kd^ was significantly blunted in 2.8 mM and 16.7 mM glucose, and trended toward reduction at 5.6 mM (Fig 1G). Likewise, after *SIX3*^kd^, there was significant blunting of insulin secretion in 2.8 mM, 5.6 mM, 16.7 mM and 25 mM glucose (Fig. 1I). GSIS data were normalized to total pseudoislet insulin content, which was not significantly altered after *SIX2*^kd^ (Fig. 1H) or *SIX3*^kd^ (Fig. 1J). This suggests that reduced insulin release from islets after *SIX2*^kd^ or *SIX3*^kd^ reflects impaired secretion. Together, our findings provide index evidence that reduced *SIX2* or *SIX3* function impairs human adult islet β-cell function.

### Elucidating the SIX2-dependent adult β-cell transcriptome

SIX2 is an established transcriptional regulator (Kobayashi et al. 2008; He et al. 2010; Self et al. 2009; Park et al. 2012) but SIX2-dependent genetic targets in adult islet β-cells are unknown. To identify genes regulated by SIX2, we purified β-cells from pseudoislets after *SIX2*^kd^ (Fig. 2A-C; Methods). Intracellular labeling with antibodies against insulin and glucagon followed by flow cytometry and GFP gating (Fig. 2C and Supplemental Fig. 2A, Peiris et al. 2018) enriched for INS^+^ GFP^+^ β-cells; qRT-PCR analysis of this cell subset confirmed enrichment of mRNA encoding INS, and depletion of GCG, or of the acinar and ductal cell markers, CPA1 and KRT19 (Supplemental Fig. 2B). We verified efficient shRNA-mediated suppression of *SIX2* in INS^+^ GFP^+^ β-cells compared to control INS^+^ GFP^+^ β-cells (Supplemental Fig. 2C). We then produced and sequenced RNA-Seq libraries from *SIX2^kd^* and control β-cells (n=4 independent donors; Methods). Pearson correlation and hierarchical clustering analysis revealed clustering of *SIX2*^kd^ and control samples from the same donor (Supplemental Fig. 2D and E), reflecting the expected inter-donor variability we and others have previously reported (Arda et al. 2016; Peiris et al. 2018; Enge et al. 2017; Segerstolpe et al. 2016). We used the DE-Seq2 algorithm (Love et al. 2014) to identify differentially expressed (DE) genes following *SIX2*^kd^. Expression of 1242 genes, including *SIX2* itself, was significantly decreased after *SIX2*^kd^ (Fig. 2D,G: Supplemental Table 2), while expression of 928 genes was significantly increased (*P*<0.05; Fig. 2I: Supplemental Table 3). By contrast, we did not detect changes in expression of β-cell *SIX3* upon *SIX2^kd^* (Fig. 2E). Gene ontology (GO) term analysis (Fig. 2F) suggested unifying molecular functions in these enriched gene sets, including terms related to adult β-cell function, proliferation and cell cycle regulation.

**Figure 2.**
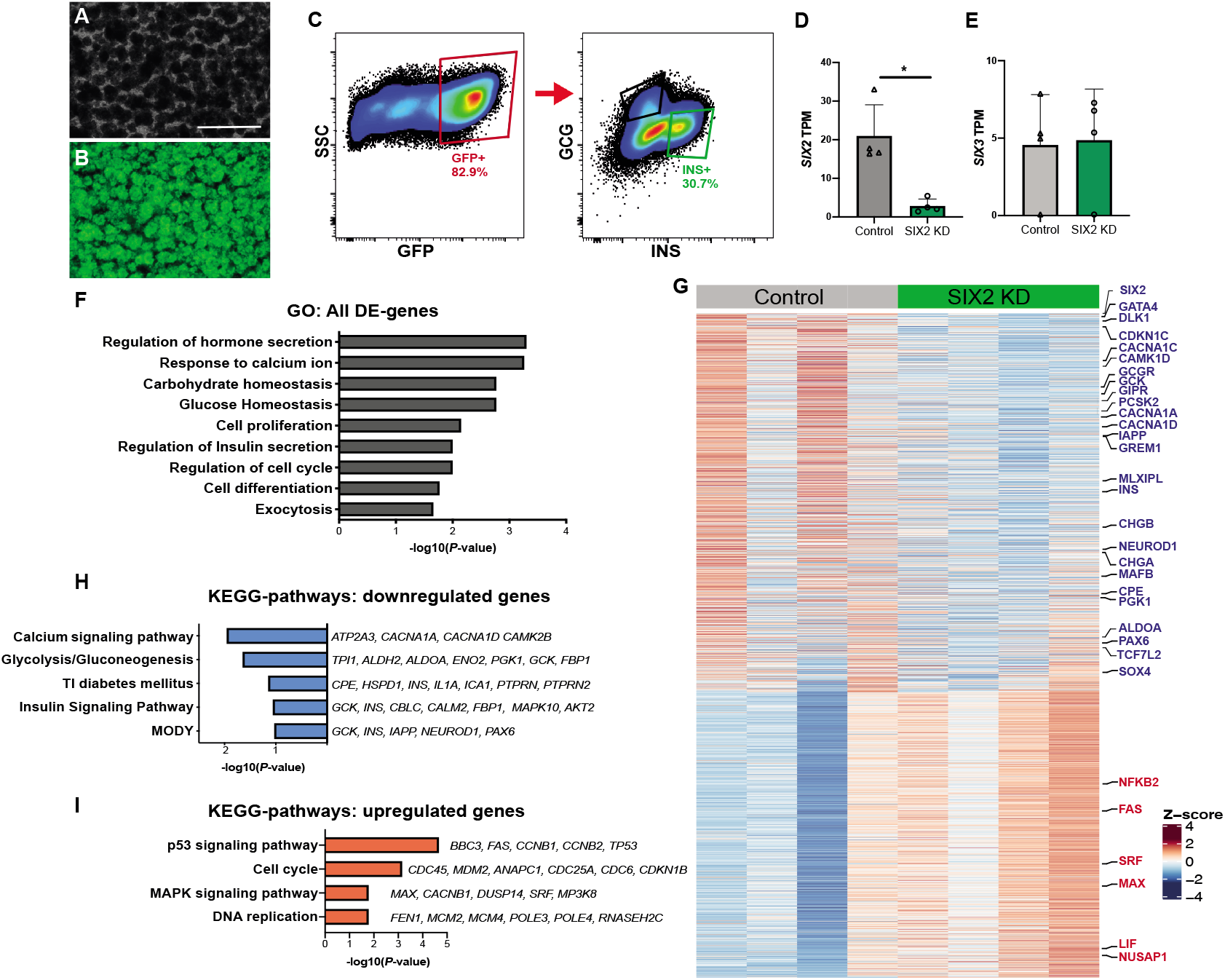
RNA-Seq of *SIX2*^kd^ β-cells reveals genes regulated by SIX2 in primary human islets. (A,B) *SIX2^kd^* human pseudoislets: (A) bright field; (B) blue light (488 nm), scale bars = 500 μm. (C) FACS scheme used to sort GFP+ β-cells. (D-E) Normalized transcript levels of (D) *SIX2*, (E) *SIX3* in GFP expressing β-cells, control (grey bar) or *SIX2^kd^* (green bar) (n=4). (F) GO term enrichment in genes de-regulated in β-cells post-*SIX2*^kd^. (G) Heatmap showing all differentially expressed (DE) genes in β-cells post-*SIX2*^kd^. (H-I) KEGG pathway enrichment in genes (H) downregulated or (I) upregulated in β-cells post-*SIX2*^kd^ (*P*<0.1; n=4 independent donors). The data is presented as mean, error bars represent the standard error. * P<0.05

Genes with decreased expression in *SIX2*^kd^ β-cells included those encoding cardinal β-cell factors like *INS*, *CHGA, CHGB* and *IAPP*, insulin processing enzymes like *CPE* and *PCSK2*, and transcription factors like *PAX6, NEUROD1, NKX6.1, MLXIPL, TCF7L2, ESRRG* and *MAFB* (Fig. 2G-H: Supplemental Table 2). In addition, we noted severe reduction of mRNAs encoding glucokinase (*GCK*), the principal sensor of glucose flux in β-cells (Matschinsky et al. 1993), Glucagon receptor (*GCGR*), regulators of glycolysis and β-cell stimulus-secretion coupling encoded by *TPI1, ALDH2, ALDOA, ENO2, PGK1* and *FBP1*, and *CAMK1D*, a postulated type 2 diabetes risk gene (Thurner et al. 2018; Miguel-Escalada et al. 2019) (Fig. 2G). In contrast, we observed increased levels of mRNAs encoding regulators of DNA replication and cell cycle factors in *SIX2^kd^* β-cells, including *NUSAP1, MAX* and *SRF* (Fig. 2G,I). This is consistent with prior findings (Arda et al. 2016) providing evidence that SIX2 expression may enforce β-cell cycle arrest, and studies linking β-cell cycle exit to enhanced function (Helman et al. 2016). GO and KEGG pathway analysis of DE genes (Methods) revealed significant enrichment of terms including regulation of hormone secretion, glucose homeostasis, calcium signaling, and insulin signaling (Fig. 2 F,H-I). Thus, these data suggest that *SIX2* is required to maintain hallmark adult β-cell functions involved with insulin production and processing, glucose sensing, and proliferation, and support our finding of impaired GSIS after *SIX2^kd^* in adult human islets.

To validate our findings further, we assessed if genes regulated by *SIX2^kd^* were enriched in gene sets whose expression changed in β-cells expressing *SIX2* (Arda et al. 2016; Blodgett et al. 2015) (see Methods). For example, we asked if the set of genes with *decreased* expression after *SIX2*^kd^ overlapped with DE genes whose mRNA normally *increased* with advancing age. In this case, we identified 287 genes with decreased expression after *SIX2*^kd^ in β-cells, whose expression was significantly increased in SIX2^+^ adult β-cells compared to fetal or juvenile SIX2^neg^ β-cells (*P*< 3.873e-30; Fig. 3A-B; Supplemental Table 7). These included *CDKN1C* and *CDKN1A*, which encode inhibitors of cell proliferation, *CACNA1C* and *CACNA1D*, which encode calcium channels, *GCGR*, which encodes a receptor for glucagon and the incretin GLP-1 (Svendsen et al. 2018), *MLXIPL* (also known as ChREBP), a glucose-activated transcription factor that regulates *GCGR* (Iizuka et al. 2012), *GREM1, IAPP* and *DLK1* (Fig. 3A-B; Supplemental Table 7). Based on similar logic, we identified 146 genes with significantly *increased* expression after *SIX2*^kd^, whose expression is normally *decreased* in SIX2^+^ adult β-cells (Arda et al. 2016; Blodgett et al. 2015) (*P*<5.2e-5; Fig. 3B; Supplemental Table 7). These included *NUSAP1* and *SRF*, postulated regulators of islet cell proliferation (Zeng et al. 2017), and *RFX7*, a marker of pancreatic progenitor cells (Kim-Muller et al. 2016). We also compared this dataset to DE transcriptomic data recently reported after *SIX2* loss in β-like cells (“SC-β”) generated from a human embryonic stem cell line (Velazco-Cruz et al. 2020). We found a general lack of concordance in DE genes (Methods): specifically, we observed 30% (377/1242) overlap of genes with reduced expression and 14% (132/928) overlap of genes with increased expression after *SIX2*^kd^ (Supplemental Fig. 3A-B, Supplemental Table 12). Many mRNAs that decreased after *SIX2^kd^* in native β cells, like *MAFB, GIPR* and *GREM1*, did not change after *SIX2*^kd^ in SC-β cells. Other mRNAs that decreased after *SIX2*^kd^ in adult β cells, like *CDKN1C, GCGR, TCF7L2* and *NKX6.1*, were found *increased* after *SIX2^kd^* in SC-β cells (Velazco-Cruz et al. 2020). These findings indicate that genetic programs in native human β cells and SC-β cells are distinct and demonstrate advantages of investigating *SIX2*-dependent gene expression in primary adult β-cells.

**Figure 3.**
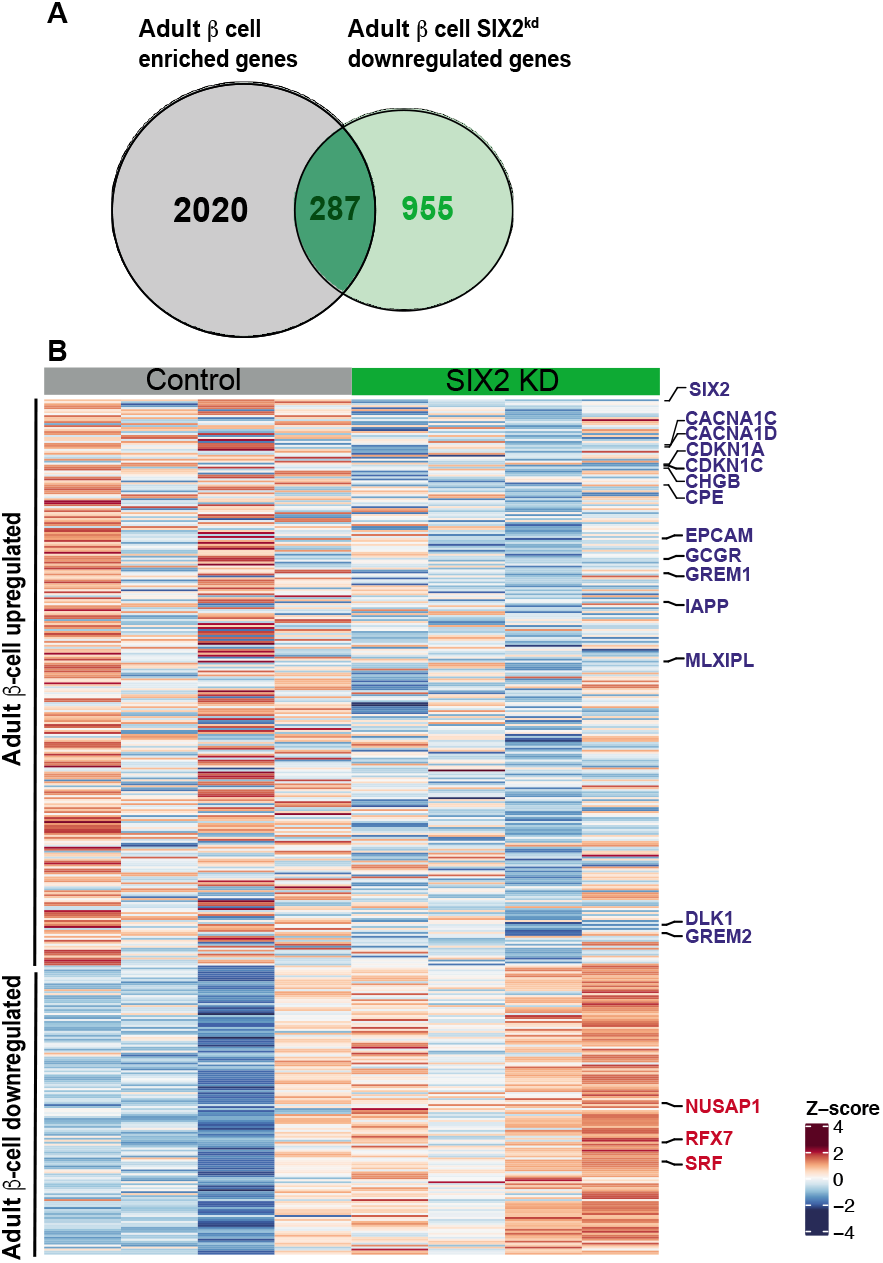
*SIX2^kd^* in primary β-cells results in downregulation of genes enriched in adult β-cells. (A) Venn diagram showing adult β-cell genes downregulated post-*SIX2*^kd^. (B) Heatmap of adult-downregulated orjuvenile-upregulated genes in adult β-cells post-*SIX2*^kd^.

### Identifying direct genetic targets of SIX2 regulation in human β-cells

To identify direct genetic targets of SIX2 in primary human islet cells, we performed cleavage under targets and release using nuclease (CUT&RUN, Skene and Henikoff 2017; Hainer et al. 2019), which enables the sensitive detection of genomic loci bound by TFs (Fig. 4A: Methods). Because antibodies that detected native islet SIX2 for CUT&RUN were not available, we mis-expressed in pseudoislets a transgene encoding human SIX2 tagged with the FLAG immuno-epitope (*SIX2-FLAG*) from a rat insulin promoter element (RIP, Karlsson et al. 1987), then sequenced DNA bound by the SIX2-FLAG protein with an anti-FLAG antibody (Supplemental Fig. 4A). Thus, interpretation of CUT&RUN here is qualified by the possibility that SIX2-FLAG may bind sites not bound by native SIX2. We used the HOMER algorithm (Heinz et al. 2010) to identify genomic regions that were bound by SIX2-FLAG in samples from three islet donors (*P*< 0.01 compared to IgG controls). Heatmap visualization of independent peaks (Fig. 4B) as well as histogram plotting of averaged reads (Supplemental Fig. 4B) showed enrichment of read densities in the peak centers for the SIX2-FLAG libraries, whereas IgG controls showed minimal enrichment at these sites (Fig. 4B, Supplemental Fig. 4B). As further validation of the specificity of CUT&RUN, we found that SIX2-FLAG-bound genomic peaks were significantly enriched for the SIX2 DNA-binding motif, as well as the SIX1 DNA-binding motif, and motifs of other β-cell-enriched TFs, like MAFB (Fig. 4C). We used the GREAT algorithm (McLean et al. 2010) to associate SIX2-FLAG-bound genomic regions to 10270 genes (Fig. 4D). Inadequate yields precluded analogous SIX3-FLAG studies (Methods).

**Figure 4.**
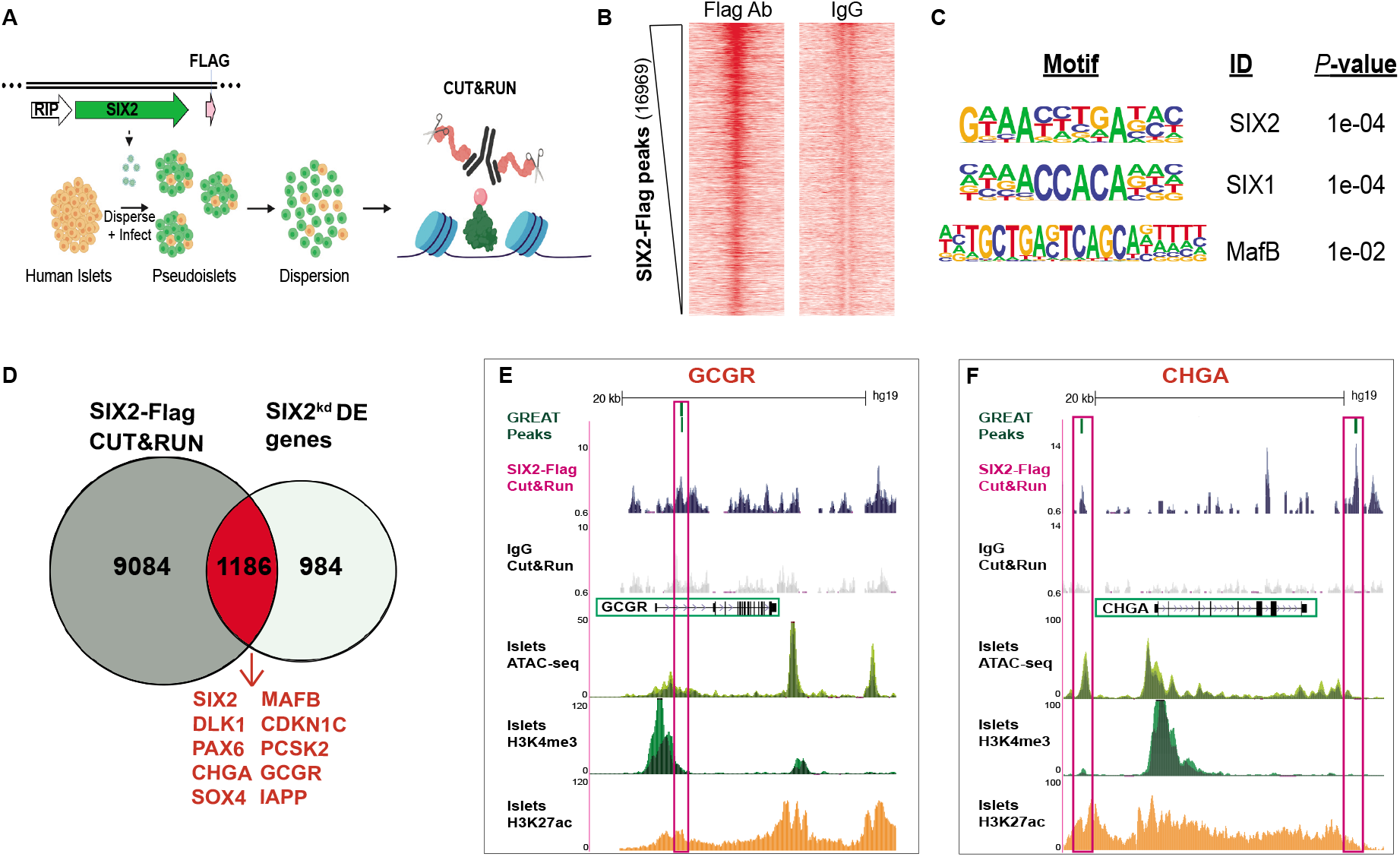
Identification of presumptive SIX2 direct genetic targets in primary human β-cells using CUT&RUN. (A) Schematic detailing the CUT&RUN approach: Pseudoislets overexpressing SIX2-Flag under the RIP promoter were used for CUT&RUN with anti-Flag antibody (n=3 independent donors). (B) Heatmap showing enrichment of peaks read densities at the center of the peak for the CUT&RUN libraries generated with Flag antibody, but not for IgG. Peaks were called using HOMER. (C) Enriched MOTIFs in the differential peaks were identified by HOMER. (D) Overlap of the SIX2-associated genomic regions and the *SIX2^kd^* DE genes. (E-F) Tracks showing SIX2-Flag genomic regions associated to (F) GCGR or (G) CHGA. Accessible chromatin regions in human islets are shown by ATAC-seq, H3K4me3 and H3K27ac ChiP-seq tracks. SIX2-Flag CUT&RUN peaks are shown in pink boxes (note for GCGR two peaks are shown), and regulated genes highlighted in green boxes.

To nominate candidate genes that might be directly regulated by SIX2, we then identified genes that (1) neighbored SIX2-FLAG binding sites and (2) had differential expression after *SIX2*^kd^. After intersection of CUT&RUN with 2170 DE genes after *SIX2*^kd^, we identified 1186 ‘overlapping genes’ (Fig. 4D; Supplemental Table 4). 64% of these genes (754/1186) had reduced mRNA levels after *SIX2*^kd^, consistent with a role for SIX2 in activating β-cell gene expression. Presumptive direct SIX2 targets identified by this approach included *PAX6, IAPP, MAFB, CDKN1C, DLK1, PCSK2, GCGR, MLXIPL* and *CHGA*. As expected, SIX2-FLAG-bound genomic regions in presumptive SIX2 target genes were found in accessible chromatin and co-localized with activation-associated H3K4me3 and H3K27ac histone marks identified by prior islet ATAC-seq and ChIP-seq studies (Mularoni et al. 2017). This alignment further supports the conclusion that SIX2 binds active genomic regulatory elements governing hallmark β-cell genes (Fig. 4E-F; Supplemental Fig. 4C-E; Supplemental Table 4). Thus, our targeted nuclease-based analysis revealed hundreds of SIX2-associated candidate regulatory elements, including many that likely regulate expression of hallmark β-cell factors.

### SIX3 and SIX2 regulate distinct gene sets in adult human β-cells

Gain-of-function studies have linked *SIX3* to adult human β-cell functional maturation (Arda et al. 2016). To identify genes whose expression is regulated by SIX3 in human β-cells, we used flow cytometry to purify β-cells from human pseudoislets after *SIX3*^kd^ (Supplemental Fig. 5A-C; see Methods). Like in our *SIX2*^kd^ studies, we confirmed enrichment of INS^+^ GFP^+^ β-cells - and depletion of non-β cells - with qRT-PCR analysis (Supplemental Fig. 5D), then generated, sequenced and analyzed RNA-Seq libraries (n=3 independent donors). We achieved 50% reduction of *SIX3* mRNA in purified INS^+^ GFP^+^ β-cells (Fig. 5A, Supplemental Fig. 5E). By contrast, *SIX2* mRNA levels were not detectably changed (Fig. 5A). As expected, analysis of RNA-Seq libraries with Pearson correlation analysis and hierarchical clustering revealed close clustering of *SIX3*^kd^ and control β-cell samples from the same donor (Fig. 5B, Supplemental Fig. 5F). With the DE-Seq2 algorithm (Methods), we identified 263 genes with significantly decreased mRNA levels (*P*<0.05; Supplemental Table 5), including *SIX3* itself (Fig. 5A), and 372 genes with significantly increased mRNA levels in *SIX3*^kd^ β-cells (*P*<0.05; Fig. 5E; Supplemental Table 6). Thus, the number of DE genes with increased mRNA outnumbered those with decreased mRNA after *SIX3*^kd^, unlike DE genes after *SIX2*^kd^. Gene ontology (GO) analysis suggested molecular functions of the enriched gene sets, including regulation of DNA binding, response to glucose stimulus, and DNA replication (Fig. 5C; Supplemental Fig. 5G). Upregulated DE genes associated with the latter term included established regulators of β-cell replication, like *MYC, MAX* and *INSM1* (Fig. 5C,E). Pathway analysis of the DE upregulated genes (Methods) included TGF-β signaling, Type 2 diabetes mellitus and calcium signaling pathway (Fig. 5D).

The DE genes after *SIX3*^kd^ were largely distinct from DE genes after *SIX2*^kd^; only 133/2805 DE genes (<5%) overlapped between *SIX2^kd^* and *SIX3^kd^* (Fig. 5F; Supplemental Table 8; Supplemenatl Fig. 6A-B). Moreover, among these 133 DE genes, 112/133 (84%) changed in the *opposite* direction after *SIX2^kd^* compared to *SIX3*^kd^, such as *KCNMB2, GREM2, OLIG1* and *TLE2*. After *SIX3^kd^* there was also a significant increase of mRNAs encoding genes not usually expressed in adult human β-cells. This included genes encoding factors enriched or exclusively expressed in islet a-cells or ε-cells, like *DPP4, NPNT, TMEM236, ADORA2A, GHRL*, which encodes the ε-cell hormone ghrelin, and *MBOAT4* (which encodes an acetyl-transferase for ghrelin) (Fig. 5E,G). While not reaching statistical significance, we also detected an average 2.5 fold increase of mRNA encoding GCG in *SIX3*^kd^ β-cells (Fig. 5G). In *SIX3*^kd^ β-cells, we also identified increased average levels of 70 mRNAs that are typically expressed highly in fetal β-cells, but attenuated or extinguished in adult β-cells (Arda et al. 2016; Blodgett et al. 2015) (Fig. 6A-B; Supplemental Table 9). This latter group of genes included *MYC* (Puri et al. 2018) and a ‘disallowed’ gene (Pullen et al. 2010) that encodes hexokinase (*HK2*: reviewed in Lemaire et al. 2016). Moreover, none of these mRNAs increased in β-cells after *SIX2*^kd^. Together, our findings support the view that *SIX2* and *SIX3* regulate distinct gene sets in human β-cells.

**Figure 5.**
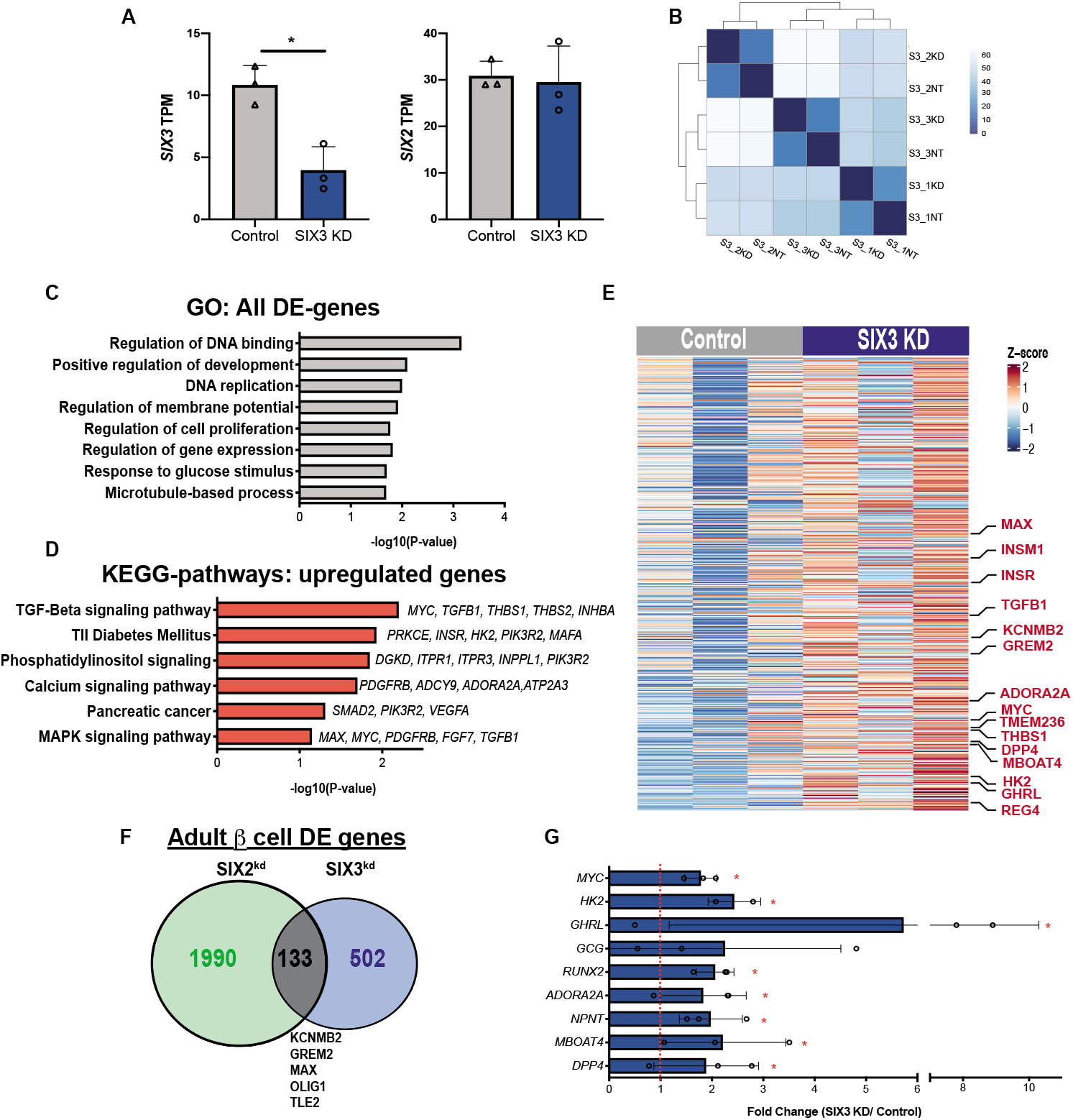
RNA-Seq of *SIX3*^kd^ β-cells reveals a distinct gene set regulated by SIX3. (A) Normalized transcript levels of *SIX3* and *SIX2* in GFP expressing β-cells, control (grey bar) or *SIX3*^kd^ (blue bar) in primaryhumanislets(n=3 independent donors). (B) Heatmap of the sample-to-sampledistances for all the samples used in this experiment. (C)GO termenrichmentin genes de-regulated in β-cells post-*SIX3*^kd^. (D) KEGGpathways enriched in genes upregulated in β-cells post-*SIX3*^kd^ (n=3independentdonors). (E) Heatmap showing all upregulated genes upon *SIX3*^kd^ in β-cells.(F) Overlapped DE genes in adult β-cells post *SIX2*^kd^ and *SIX3*^kd^.(G)Fold transcript levels of non-β-cell genes significantly alteredin β-cellspost-*SIX3*^kd^ (n = 3 independent donors). The data is presented asmean,error bars represent the standard error. * *P*<0.05

**Figure 6.**
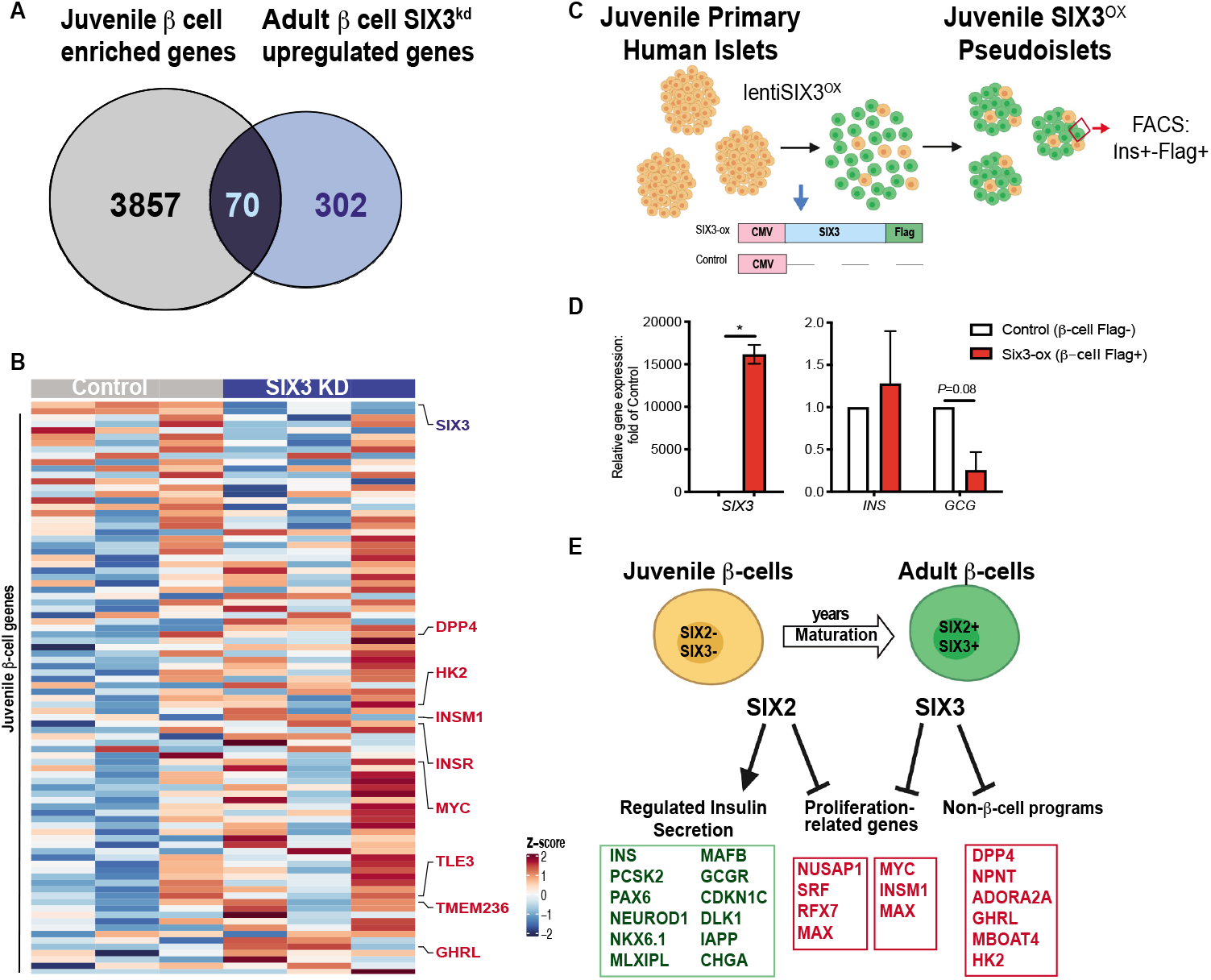
SIX3 represses non-β-cell programs in the adult β-cell. (A) Venn diagram showing juvenile β-cell genes upre-gulated in adult β-cells post-*SIX3*^kd^. (B) Heatmap of adult-downregulated or juvenile-upregulated genes in adult β-cells post-*SIX3*^kd^. (C) Schematic detailing the pseudoislet technique used to overexpress SIX3-Flag (SIX3-ox) in juvenile pseudoislets (n=2 independent donors), and of the constructs used to overexpress *SIX3* in juvenile pseudoislets: FACS was used to sort Flag+ β-cells. (D) *SIX3, INS* and *GCG* mRNA expression in Flag+ juvenile β-cells post FACS, control (white bar) or SIX3-ox (red bar) (n=2 donors). (E) Schematics showing proposed coordinated regulation of maturation by SIX2 and SIX3 in the β-cell. The data is presented as mean, error bars represent the standard error. * P<0.05

We previously showed that *SIX3* expression in pseudoislets from juvenile human donors aged 0.5 to 2 years (which lack *SIX3* expression) was sufficient to stimulate insulin secretion *in vitro* (Arda et al. 2016); however, transcriptome studies were not performed. To assess changes of gene expression stimulated by *SIX3*, we mis-expressed a *SIX3-FLAG* transgene in pseudoislets from two juvenile donors (ages 1.5 and 3 years: Fig. 6C; Supplemental Table 1). Flow cytometry and western blotting verified and quantified expression of transgenic *SIX3-FLAG* (Supplemental Fig. 7A-C). qRT-PCR analysis of purified β-cells showed a reduction in *GCG* levels and an average increase of *INS* mRNA levels following *SIX3* misexpression (Fig. 6D), as previously reported (Arda et al. 2016). RNA-Seq of purified SIX3-FLAG^+^ INS^+^ β-cells (Methods) confirmed *reduced* mRNAs encoding GCG, HK2, MYC, TMEM236, MBOAT4, RUNX2, GREM2 and NPNT, mRNAs found *increased* after *SIX3*^kd^ in primary β-cells (Supplemental Fig. 7D; Supplemental Table 10). Thus, *SIX3* gain- and loss-of-function studies here produced *reciprocal* changes in expression of multiple genes, supporting the view that SIX3 suppresses adult β-cell expression of gene sets expressed abundantly in fetal or neonatal β-cells, or in adult a- and ε-cells. We conclude that SIX3 re-enforces mature β-cell function, in part, by suppressing fetal gene expression programs and alternative islet cell fates (Fig. 6E).

### Human β-cell expression of SIX2 is reduced in islets from T2D donors

It remains unknown whether the expression of *SIX2* or *SIX3* changes in islet β-cells obtained from T2D patients. We found that both *SIX2* and *SIX3* mRNA appeared to be reduced in purified *whole islets* from subjects with established T2D (n=4), compared to islets from non-diabetic controls (n=7; Fig. 7A: Donor information in Supplemental Table 1). By contrast, expression of *GPD2* and *LEPROTL2*, previously found upregulated in prior studies (Segerstolpe et al. 2016), was unaltered in T2D islets (Fig. 7A).

**Figure 7.**
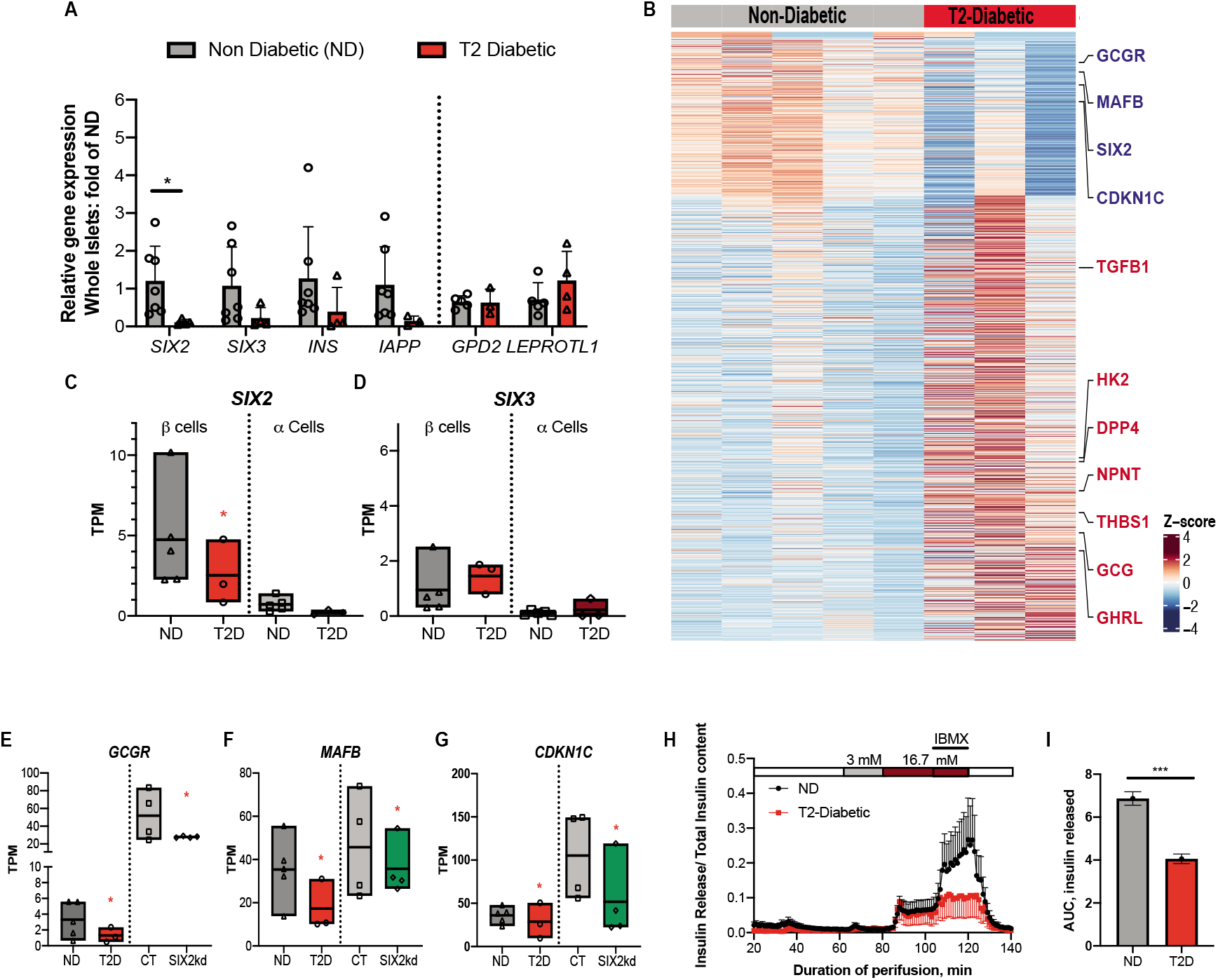
*SIX2* expression is reduced in β-cells from Type 2 Diabetic donors. (A) Gene expression levels in whole islets from non-diabetic (ND, grey bars) (n=7) or type 2 diabetic (T2D, red bar) (n=4) donors. (B) Heatmap showing DE genes in ND versus T2D β-cells. (C-D) Boxplots displaying TPM of (C) *SIX2* and (D) *SIX3* in ND β-cells (dark-grey bars) and a-cells (light-grey bars) (n=5 donors) or T2D β-cells (red bars) and a-cells (dark-red bars) (n=3 donors). (E-G) Boxplots displaying TPM counts of (E) *GCGR*, (F) *MAFB* and (G) *CDKN1C* in β-cells of ND (n=5 donors) and T2D (n=3 donors); the expression of which is also DE in adult β-cells post-*SIX2*^kd^, control (grey bar), *SIX2^kd^* (green bars). (H) Insulin secretion of ND (n=5 donors) versus T2D islets (n=3 donors): see Methods. (I) Plot of the total area under the curve of the released insulin. The data is presented as mean, error bars represent the standard error. Box plots show the mean. Red *, *P*<0.01; Black *, *P*<0.05, *** *P*<0.001

Bulk RNA-Seq from FACS-purified a- and β-cells was used to assess *SIX2* and *SIX3* expression in β-cells (Fig. 7B-D; see Methods). Transcriptome analysis confirmed significant reduction of *SIX2* mRNA in purified β-cells from T2D islets, compared to β-cells from control islets (*P* < 0.01: Fig. 7B-C; Supplementary Table 11), while *SIX3* mRNA levels were not significantly changed (Fig. 7D). Thus, *SIX2* expression was reduced in β-cells in islets from cadaveric T2D donors. Moreover, β-cell regulation of *SIX2* and *SIX3* in these T2D islets was uncoupled. We also detected little to no expression of *SIX2* and *SIX3* in purified a-cells from control or T2D islets (Fig. 7C-D). In addition to reduced *SIX2* expression in β-cells, we also noted impaired expression of *SIX2* target genes, including *MAFB, GREM1, GCGR*, and *CDKN1C*, a candidate T2D risk gene (Fig. 7B,E-G, Supplemental Table 11). Consistent with our studies of impaired insulin secretion after *SIX2^kd^* (Fig. 1G), glucose-stimulated insulin secretion studies revealed impaired insulin secretion by islets from subjects with T2D compared to non-diabetic controls (Fig. 7H-I: see Methods).

Although SIX3 mRNA levels were not detectably changed in our sampling of T2D β cells, a subset of *SIX3* targets was differentially expressed in T2D β cells, including *DPP4, NPNT* and *GHRL*; TGF-β signaling factors encoded by *TGFB1* and *THBS1*; and the disallowed gene *HK2* (Fig. 7B). This raises the possibility that factors collaborating with SIX3 to regulate these genes might be changed in T2D β-cells. Together, these findings suggest that dysregulation of *SIX2*- and *SIX3*-dependent genetic programs could contribute to impaired islet β-cell fate and function in T2D. These findings also support prior genomewide association studies linking the locus encoding *SIX2* and *SIX3* to risk for T2D and diabetes-related traits (Spracklen et al. 2018; Kim et al. 2011; Spracklen et al. 2020; Hachiya et al. 2017; Varshney et al. 2017), raising the possibility that genetic influences might additionally modulate β-cell expression of *SIX2* or *SIX3*.

## Discussion

Here we overcame inherent challenges facing postnatal human developmental studies to investigate the roles of SIX2 and SIX3 in human pancreatic β-cell maturation. Using genetic approaches, we show that SIX2 is required for expression of multiple hallmark genes in human β-cells. In addition to regulation of genes governing β-cell function, we show that SIX3 - unlike SIX2 - suppresses expression of genes typically expressed in a-cells or other non-β-cells. Thus, our studies provide index evidence that SIX2 and SIX3 regulate distinct sets of genetic targets in adult human β-cells. shRNA-mediated suppression of either *SIX2* or *SIX3* expression in primary human islets impaired regulated insulin secretion by β-cells. Supporting these findings, we found evidence of reduced expression of *SIX2* and downstream targets in islet β-cells from human subjects with T2D, which coincided with significantly reduced insulin secretion by these islets. In sum, our study unveils a requirement for *SIX2* and *SIX3* in establishing and maintaining adult human β-cell function and fate (Fig. 6E).

Prior to our study, it was unclear what roles SIX2 and SIX3 had in adult human β-cell function. *SIX2* and *SIX3* are co-expressed in adult human β-cells, and developmental studies of human islet cells have revealed coincident increases of *SIX2* and *SIX3* expression after the first decade of life (Arda et al. 2016; Arda et al. 2018; Blodgett et al. 2015). Moreover, studies of putative *SIX2* and *SIX3 cis*-regulatory elements in humans and other systems have suggested these genes may be co-regulated (Spracklen et al. 2018; Suh et al. 2010). A common set of SIX2 and SIX3 targets identified here includes regulators of cell cycle progression. However, the onset of *SIX2* and *SIX3* in human β-cells occurs well after the period of neonatal expansion (Arda et al. 2016; Blodgett et al. 2015) suggesting that the post-mitotic state of β-cells is established by other factors, and then re-enforced by SIX2 and SIX3. Consistent with this possibility, we observed that reduction of *SIX2* alone or *SIX3* alone did not increase markers of β-cell S-phase like *MKI67* (Supplementary Tables 2-5). Moreover, we showed in our prior work that mis-expression of either SIX2 or SIX3 was sufficient to suppress proliferation of the human β-cell line EndoCβH1 (Arda et al. 2016). Enforcement of the post-mitotic state in β-cells has been linked to attainment of mature function (Helman et al. 2016; Puri et al. 2018; Mandelbaum et al. 2019).

The majority of differentially-expressed genes after *SIX2*^kd^ showed reduced expression, including those encoding crucial human β-cell factors like Insulin and Glucokinase, and essential TFs that coordinate pancreatic islet development and β-cell function in humans. Consistent with this, we observed reduced Insulin secretion after *SIX2*^kd^. By contrast, after *SIX3*^kd^ the majority of DE genes showed increased expression. These included transcripts encoding factors not normally expressed in healthy β-cells, like the e-cell hormone Ghrelin, the a-cell-enriched protease DPP4, and the disallowed factor HK2 (Dhawan et al. 2015). These *SIX3*^kd^ findings are consistent with prior studies suggesting that SIX3 can function as a transcriptional repressor (Kobayashi et al. 2001). Thus, our loss-of-function studies revealed SIX2- and SIX3-dependent mechanisms that regulate native maturation and fate of human β cells. Here, the degree of SIX2 mRNA reduction after *SIX2^kd^* was greater than the degree of SIX3 mRNA loss after *SIX3*^kd^; thus, future studies that achieve more complete SIX2 or SIX3 loss of function could identify additional β-cell genetic targets. Our study was also limited by the inherent variability of cadaveric human islet donors, as we and others have previously reported (Arda et al. 2016; Peiris et al. 2018; Enge et al. 2017; Segerstolpe et al. 2016). While *SIX3* expression is restricted to the β-cell, *SIX2* is also expressed in islet δ cells (Baron et al. 2016; Muraro et al. 2016). Thus, changes observed after *SIX2^kd^* could reflect both β-cell autonomous and non-autonomous mechanisms. Studies here also revealed that *SIX2* and *SIX3* expression in β-cells can be genetically uncoupled and correspond well with our data showing that *SIX2* and *SIX3* regulate distinct β-cell gene sets. Thus, distinct mechanisms likely govern expression and activity of SIX2 and SIX3 in β-cells from healthy and diabetic cadaveric donors.

Elucidating how *SIX2* and *SIX3* expression are regulated should be aided by our identification of their targets in human β-cells. Prior studies have shown that transcription factors like *SIX2* and *SIX3* expressed in human β-cells show increased expression with age (Arda et al. 2016; Aguayo-Mazzucato et al. 2011; Blodgett et al. 2015; Wang et al. 2016b). For shRNA-based studies here we used islets from donors > 22 years of age, when adult levels of SIX2 and SIX3 have been established. Intense interest in *SIX2* and *SIX3* regulation also stems from association of the locus encoding these factors to T2D and related traits like fasting hyperglycemia (Spracklen et al. 2018; Kim et al. 2011; Spracklen et al. 2020; Hachiya et al. 2017; Varshney et al. 2017). However, studies of whole islet RNA, or prior single islet cell RNA-Seq investigations by us and others (Camunas-Soler et al 2020; Segerstolpe et al. 2016) were not sufficiently sensitive to detect changes of *SIX2* or *SIX3* mRNA in β-cells isolated from donors with T2D. Studies here provide index evidence that (1) expression of *SIX2*, and a subset of *SIX2*-dependent genes like *GCGR*, are significantly reduced in β-cells from T2D donors, and (2) islets from T2D donors with reduced β-cell expression of *SIX2* had impaired insulin secretion (Fig. 7). While we did not detect changes in *SIX3* expression in T2D islets here, additional studies are required to exclude the possibility that β-cell *SIX3* dysregulation is a feature of T2D.

A recent report described phenotypes after SIX2 loss in hPSC-derived β-like cells (SC-β cells) (Velazco-Cruz et al. 2020). While 25% of these Insulin^+^ SC-β cells express *SIX2* mRNA, they lack other markers of mature β-cells like MAFB or SIX3, thus precluding studies of *SIX3* loss-of-function. After shRNA-mediated suppression of *SIX2*, Velasco-Cruz and colleagues reported reduced insulin protein content, loss of glucose-stimulated insulin secretion without effects on ‘basal’ insulin secretion at low glucose concentration (2 mM), and significant changes in expression of >10,000 genes, assessed by RNA-Seq of unsorted hESC progeny, with enrichment of gene sets related to insulin secretion and calcium signaling. This included significant increases of multiple transcripts encoding islet a cell or δ cell products, like SST, DPP4, MBOAT4, and FSTL1 (Velazco-Cruz et al. 2020). These findings support the conclusion that SIX2 is required for SC-β cells derived from hESCs to acquire some features of native human β-cells.

In our study, we assessed the effects of SIX2 or SIX3 loss in native human β-cells. After *SIX2*^kd^, we observed reduction of both basal and glucose-stimulated insulin secretion, without reduction of islet insulin content. FACS purification of Insulin^+^ β-cells (and elimination of SIX2^+^ δ cells) and RNA-Seq revealed significant changes in mRNA levels of 2100 genes, with gene set analysis revealing enrichment of terms related to proliferation, insulin secretion, calcium signaling, carbohydrate metabolism and exocytosis. While some of these gene sets overlapped with those in SC-β-cells (Velazco-Cruz et al. 2020), there was an overall lack of concordance between gene sets (Supplemental Fig. 3, Supplemental Table 12). For example, after *SIX2^kd^* we observed reduced mRNA encoding the calcium channel subunits CACNA1A and CACNA1D (“calcium signaling”), incretin receptors GIPR and GCGR (“insulin secretion”), and transcription factors with established roles in native β-cell regulation like MAFB, NEUROD1, NKX6.1 and TCF7L2; these changes were not noted in SC-β cells. Moreover, we did not observe increased expression of non-β cell markers like DPP4, MBOAT4 and FSTL1 after *SIX2*^kd^. Instead, we observed increased expression of these genes, and other non-β cell or disallowed genes after *SIX3*^kd^. These contrasts raise the possibility that SIX2 activity in SC-β-cells includes ectopic functions normally fulfilled by SIX3 or other factors. Together, our findings clarify the importance of investigating SIX2 and SIX3 functions in *bona fide* adult β-cells. While models of human β cell development, like stem cell-derived insulin^+^ cells and immortalized β-cell lines have value (Sneddon et al 2018), to date they remain fundamentally different from genuine pancreatic islet cells in gene regulation, function, proliferation, and cellular composition. Our studies further suggest that simultaneous expression of both SIX2 and SIX3 may be required to produce consummately functional replacement β-cells from renewable sources, like human stem cell lines.

In summary, this study unveils *SIX2* and *SIX3* functions crucial for postnatal human islet and β-cell development and maturation, and reveals how β-cell dysfunction might develop in diabetes. Our work demonstrates that SIX2 and SIX3 coordinately govern distinct genetic programs that increase insulin production and enhance mature β-cell physiological functions, enforce β-cell fate by suppressing alternative genetic programs and suppress proliferation. Findings here also provide a unique developmental ‘roadmap’ for achieving human β-cell replacement.

## Materials and Methods

### Human Islet Procurement

De-identified human islets were obtained from healthy, non-diabetic organ donors or Type 2 Diabetic donors procured through the Integrated Islet Distribution Network (IIDP), National Diabetes Research Institute (NDRI), International Institute for the Advancement of Medicine (IIAM) and the Alberta Diabetes Institute Islet Core. For T2D studies, data from the Human Pancreas Analysis Program (HPAP-RRID:SCR_016202) Database (https://hpap.pmacs.upenn.edu), a Human Islet Research Network (RRID:SCR_014393) consortium (UC4-DK-112217 and UC4-DK-112232) was used. See Supplemental Table 1 for details.

### Constructs and lentivirus production

Lentiviral constructs coding for shRNAs targeting human SIX3 or SIX2 were obtained from Dharmacon. plenti-CMV-SIX3-cMyc-DDK was used in juvenile islet experiments (Origine). plenti-RIP-SIX2-cMyc-DDK was generated by replacing the CMV promoter of plenti-CMV-SIX2-cMyc-DDK (Origine) with the rat insulin promoter (RIP). Lentiviruses were produced by transfection of HEK293T cells with lentiviral constructs, pMD2.G (12259; Addgene) and psPAX2 (12260; Addgene) packaging constructs. Supernatants were collected and concentrated by PEG-it (System Biosciences).

### Human pseudoislet generation and transduction

Human islets were dispersed into single cells by enzymatic digestion (Accumax, Invitrogen) and transduced with 1×10^9^ viral units/1 ml lentivirus. Transduced islet cells were cultured in ultra-low attachment well plates for 5 days prior to further analysis.

### RNA extraction and quantitative RT-PCR

RNA was isolated from whole pseudoislets using the PicoPure RNA Isolation Kit (Life Technologies). For sorted β- and a-cells, RNA was isolated using the RecoverALL isolation kit (Invitrogen by Thermo Fisher Scientific). cDNA was synthesized using the Maxima First Strand cDNA synthesis kit (Thermo Scientific) and gene expression was assessed by PCR using Taqman Gene Expression Mix (Thermo Scientific) and probes: ACTIN-B, Hs4352667_m1; SIX2, Hs00232731_m1; SIX3, Hs00193667_m1; INSULIN, Hs00355773_m1; GLUCAGON, Hs00174967_m1; CPA-1, Hs00156992_m1 and KRT19, Hs01051611_gH.

### In vitro insulin secretion assays

Batches of 25 pseudoislets were used for *in vitro* secretion assays as previously described (Peiris et al. 2018). Briefly, pseudoislets were incubated at 2.8 mM, 5.6 mM, 16.7 mM, 25 mM and 25 mM + IBMX glucose concentrations for 60 min each, and supernatants were collected. Secreted human insulin in the supernatants and pseudoislet lysates were quantified using a human insulin ELISA kit (Mercodia). Secreted insulin levels are presented as a percentage of total insulin content. Perifusion data of T2-Diabetic versus non-diabetic samples were acquired from the Human Pancreas Analysis Program (HPAP-RRID:SCR_016202).

### Immunohistochemistry

Human pseudoislets were fixed for 1 h at 4 °C and embedded in collagen (Wako Chemicals) and OCT prior to sectioning and staining as previously described (Arda et al. 2016). Primary antibodies used: Guinea pig anti-Insulin, (1:1000, DAKO, A0564), Mouse anti-Glucagon (1:500, Sigma), Mouse anti-SST (1:500). Secondary antibodies were incubated at room temperature for 2 h. Images were obtained using a Leica SP2 confocal microscope.

### Intracellular staining and FACS sorting of human islet cells

Detailed protocol can be found in Peiris et al. 2018. Briefly, pseudoislets were dispersed into single cells and stained with LIVE/DEAD™ Fixable Near-IR Dead Cell Stain Kit (Life Technologies) prior to fixation with 4% paraformaldehyde. After permeabilization, cells were stained with antibodies: Guinea pig anti-insulin (1:100, Dako) followed by anti-guinea pig-Alexa Fluor® 555 (1:100, Sigma) and mouse anti Glucagon Antibody -Alexa Fluor® 647 (1:100, Santa Cruz Biotechnology). Juvenile islet cells from SIX3-Flag pseudoislets were stained with anti-FLAG antibody-555 (1:100; Biolegend). Labeled cells were sorted on a special order 5-laser FACS Aria II (BD Biosciences) using a 100 μm nozzle, with appropriate compensation controls and doublet removal. Sorted cells were collected into low retention tubes containing 50 μL of FACS buffer.

### RNA isolation and preparation of RNA-Seq libraries

A total of 20000 sorted, fixed β-cells were used for each RNA-Seq library construction of RNA with RIN number> 7. SMART-Seq v4 Ultra Low input RNA kit (Clontech) was used to amplify cDNA which was subsequently sheared resulting in 200-500 bp fragments. RNA-Seq libraries were generated using the Low Input Library Prep Kit v2 (Clontech). Barcoded libraries were then multiplexed and sequenced as paired-end 150 bp reads on the Illumina HiSeq4000 platform. A total of 8 libraries were generated from 4 different donors used for the *SIX2^kd^* (4 libraries) and the respective control β-cells (4 libraries); while 6 libraries were generated from 3 different donor β-cells used for *SIX3*^kd^ and their respective controls.

### Bioinformatic and statistical analysis

RNA-seq analysis was performed on *SIX2^kd^* and control β-cell libraries from four donors and on *SIX3*^kd^ and control β-cell libraries from three donors. FastQC v0.11.4 was used for quality control. All libraries had over 75 million reads and barcodes were trimmed using Trimgalore_0.5.0. Reads were aligned to the human genome index (hg19) using STAR v2.6.1d (Dobin et al. 2013). Transcripts per million (TPM) were quantified using RSEM v1.3.0 (Li et al. 2011). Differentially expressed genes with fold change were detected using the DESeq2 R package (Love et al. 2014) for the two experimental conditions. The Database for Annotation, Visualization and Integrated Discovery (DAVID) v6.7 was employed (Huang et al. 2009) for gene set enrichment analysis. RNA-seq datasets of genes enriched in adult β-cells versus juvenile β-cells were obtained from (Arda et al. 2016; GEO: GSE79469) and (Blodgett et al. 2015; GEO: GSE67543). The probability of finding x overlapping genes was calculated using the hypergeometric probability formula that considers the total number of genes in the genome. RNA-seq data of sorted β- and a-cells of T2D versus non-diabetic donors (HPAP-RRID:SCR_016202) and of *SIX2*^kd^ SC-β-cells (Velazco-Cruz et al. 2020; GEO: GSE147737) were analyzed as per our data, using DESeq2 R package (Love et al. 2014).

### CUT&RUN and library preparation

600,000 re-dispersed islet cells were used as input material for each CUT&RUN, which was performed from 3 donors, as described (Skene and Henikoff 2017; Hainer et al. 2019). Briefly, nuclei were extracted with nuclear extraction buffer and added to concanavalin A beads slurries (Polysciences). After blocking, the nuclei/beads were washed in Wash Buffer and resuspended with rabbit anti-Flag (Sigma-Millipore, F7425) or IgG (Millipore) antibodies at 4C overnight. Protein A-micrococcal nuclease (pA-MN; EpiCypher donation) was added to a concentration of 1:400 to nuclei. Cleavage was induced by 100mM CaCl2 at 0° C for 30 minutes. DNA fragments were released at 37° C for 20 minutes and purified using phenol/chloroform/ isoamyl alcohol followed by chloroform extraction and precipitated with glycogen and ethanol. DNA was resuspended in 0.1X TE and used for library construction with NEBnext Ultra II library kit. Libraries were sequenced as 2×75 on HiSeq4000.

### CUT&RUN data analysis

Paired-end reads were trimmed and aligned as per CUT&RUNtools (Zhu et al. 2019). Briefly, Trimmomatic (Bolger et al. 2014) was used for trimming and Bowtie2 (Langmead et al. 2012) for alignment. HOMER (Heinz et al. 2010) was used for peak calling. Genome browser tracks were generated from mapped reads using the ‘‘makeUCSCfile’’ command. Peaks were called using the ‘‘findPeaks’’ command. The GREAT algorithm was used for gene annotation (McLean et al. 2010). Motifs were identified using the ‘‘findMotifs’’ command. *P* values for motif enrichment were performed by HOMER software, using a binomial test.

### Statistical analysis

For qRT-PCR and GSIS, the number of biological or technical replicates (n), measure of central tendency (e.g. mean), standard deviation and statistical analysis is detailed in the figure legend. Graphs and statistical analysis were produced and performed using GraphPad Prism (version 8) software.

## Data visualization

Cytometry data was analyzed and graphed using FlowJo software (TreeStar v.10.8). Heatmaps were made with ComplexHeatmap. Browser tracks were made with the UCSC genome browser. The graphics were made with BioRender.

## Data availability

The data discussed in this publication have been deposited in NCBI’s GeneExpression Omnibus (Edgar et al., 2002) and are accessible under accession number.

## Acknowledgments

We thank past and current members of the Kim group for advice and encouragement, especially Dr. Y. Hang, S. Park and A. Ibarra Urizar for technical guidance and advice, Drs. C. Chang, Y. Hang (Kim group), and R. Bottino (Allegheny Health Network) for assistance in tissue procurement, and Dr. M. Angulo (K. Chua group, Stanford) for help with pulldowns and the CUT&RUN protocol, Dr. G. Oliver (St. Judes) for initial discussions about SIX2 and SIX3 and N. Koska for help with antibody testing, and members of the Kim lab for comments on the manuscript. We thank Dr. R. Nair (Diabetes Genomics Analysis Core, SDRC) for help with bioinformatics and programming. We thank Professors K. Loh, and A. Gloyn for advice and encouragement. We gratefully acknowledge organ donors and their families, and islet procurement through the Alberta Diabetes Institute Islet Core, Integrated Islet Distribution Program (U.S. NIH UC4 DK098085), the National Disease Research Interchange, and the International Institute for the Advancement of Medicine. R.J.B. was supported by a postdoctoral fellowship from JDRF (3-PDF-2018-584-A-N) and is on leave from the Animal Biotechnology Laboratory, Facultad de Agronomía, Universidad de Buenos Aires/INPA CONICET, CABA, Argentina. H.P. was supported by fellowships from the Maternal and Child Health Research Institute (School of Medicine, Stanford University, UL1TR001085), the American Diabetes Association (1-16-PDF-086) and a Young Investigator Award from the Stanford Institute for Immunity, Transplantation and Infection, R.L.W. by fellowships from the Division of Endocrinology National Institutes of Health T32 training grant in the Dept. of Medicine, Stanford University (DK007217-41, to A. Hoffman and F. Kraemer) and JDRF (3-PDF-2020-931-A-N) and S.K. by a fellowship from the Larry L. Hillblom Foundation (2017-D-008-FEL).Work in the Kim lab was supported by NIH awards (R01 DK107507; R01 DK108817; U01 DK123743 to S.K.K.), and JDRF Northern California Center of Excellence (to S.K.K. and M. Hebrok). Work here was also supported by NIH grant P30 DK116074 (S.K.K.), and by the Stanford Islet Research Core, and Diabetes Genomics and Analysis Core of the Stanford Diabetes Research Center.

## Author contributions

R.J.B. and S.K.K. conceptualized the study and guided the work. Methodology: R.J.B., S.K.K.; Investigation: R.J.B., J.L., H.P., R.L.W., S.K., M.S.F., X.G.; Writing: R.J.B. and S.K.K. wrote the manuscript with input from all coauthors; Visualization: R.J.B.; Supervision: S.K.K.; Funding acquisition: R.J.B, R.L.W. H.P., S.K., M.S.F., S.K.K.

