## Supplemental Figures, Suppl Tables 1, 7,8,9,12 for "SIX2 and SIX3 coordinately regulate functional maturity and fate of human pancreatic β cells"

Supplemental Fig. 1

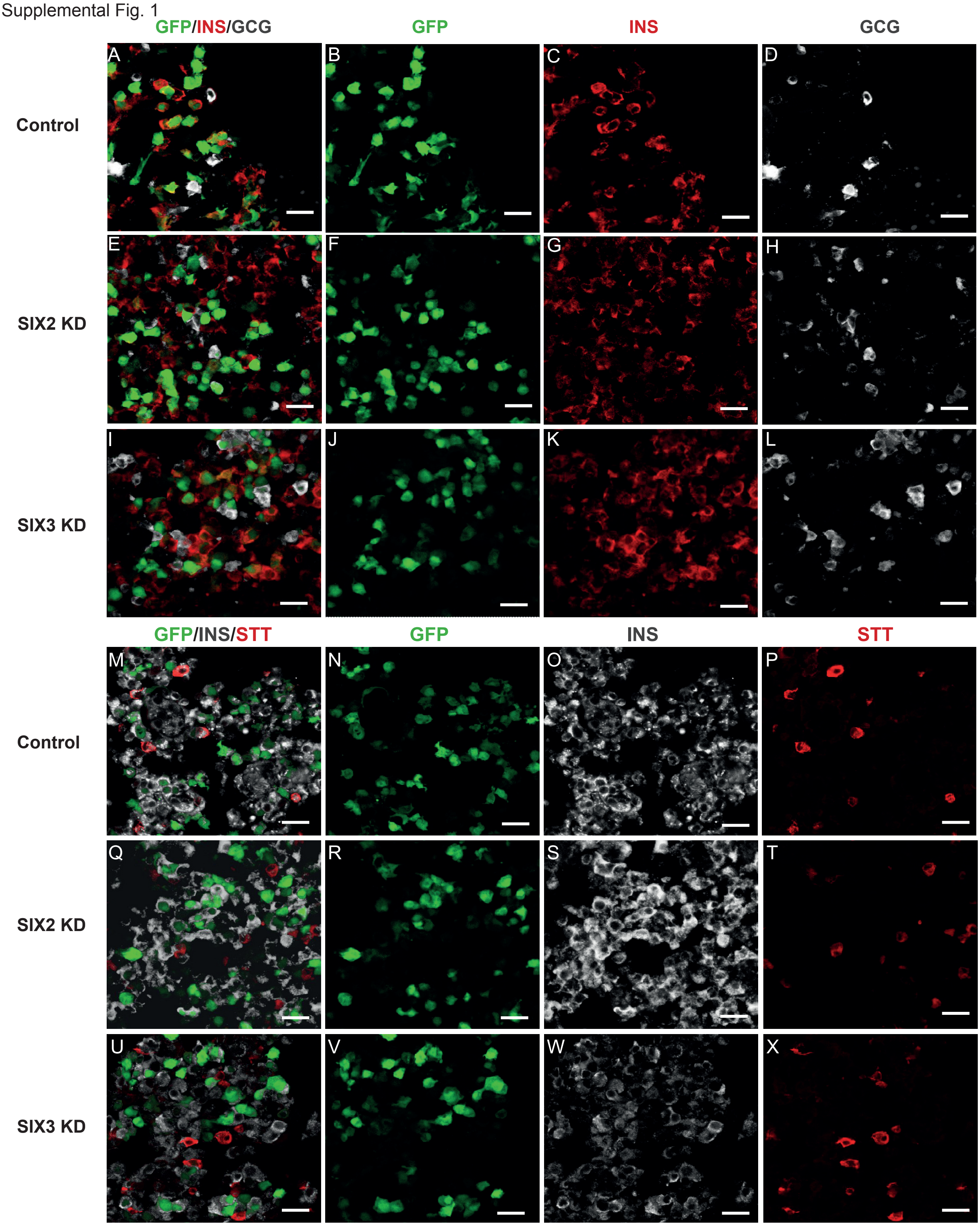

Supplemental Fig. 2

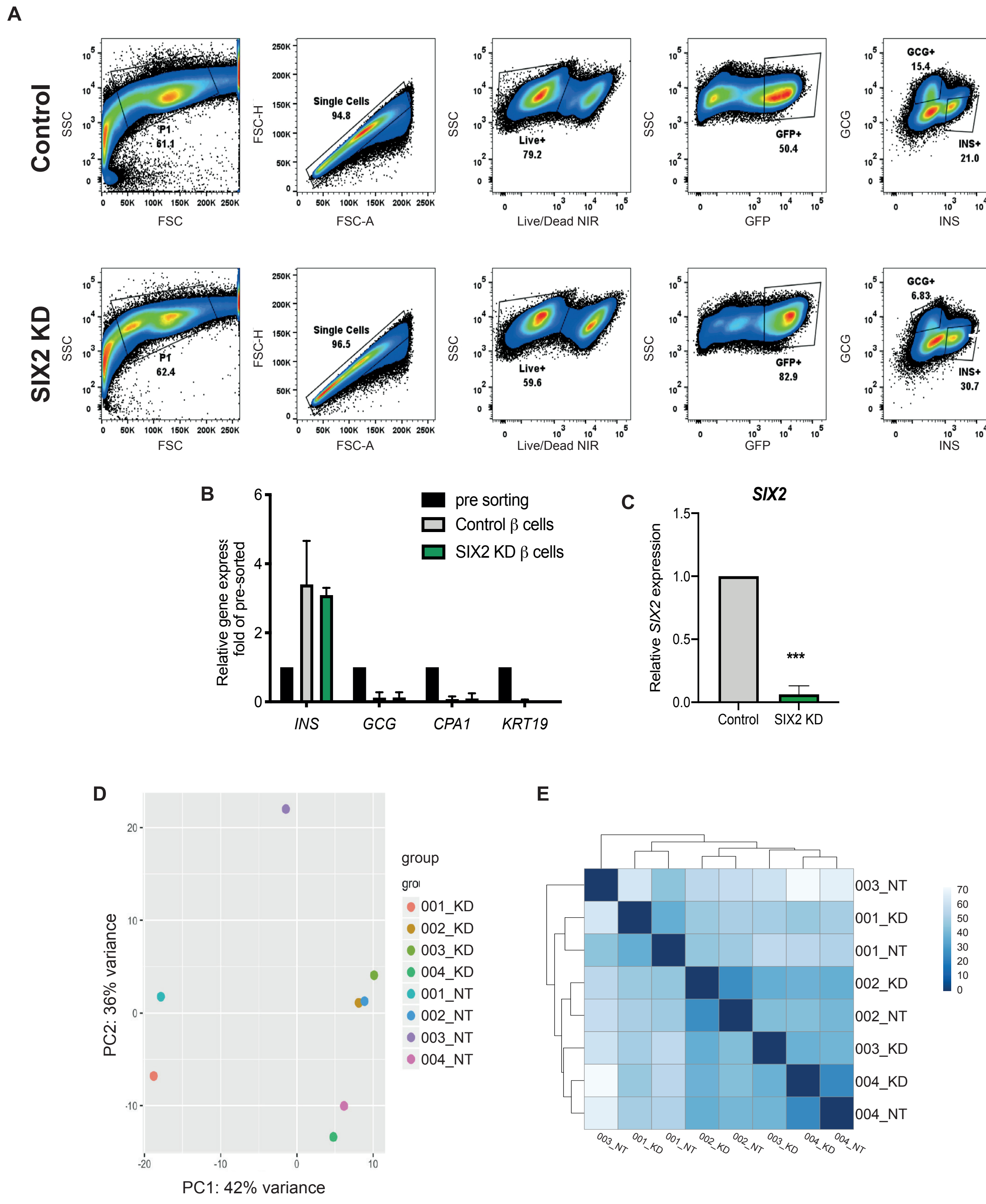

Supplemental Fig. 3

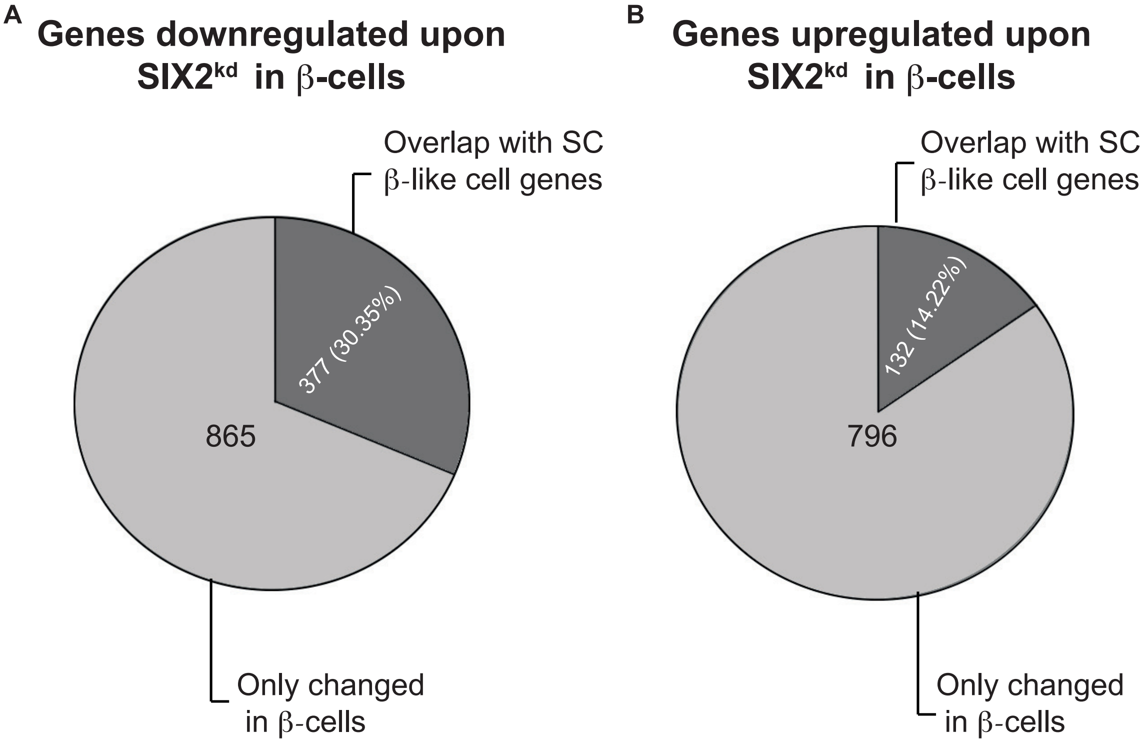

Supplemental Fig. 4

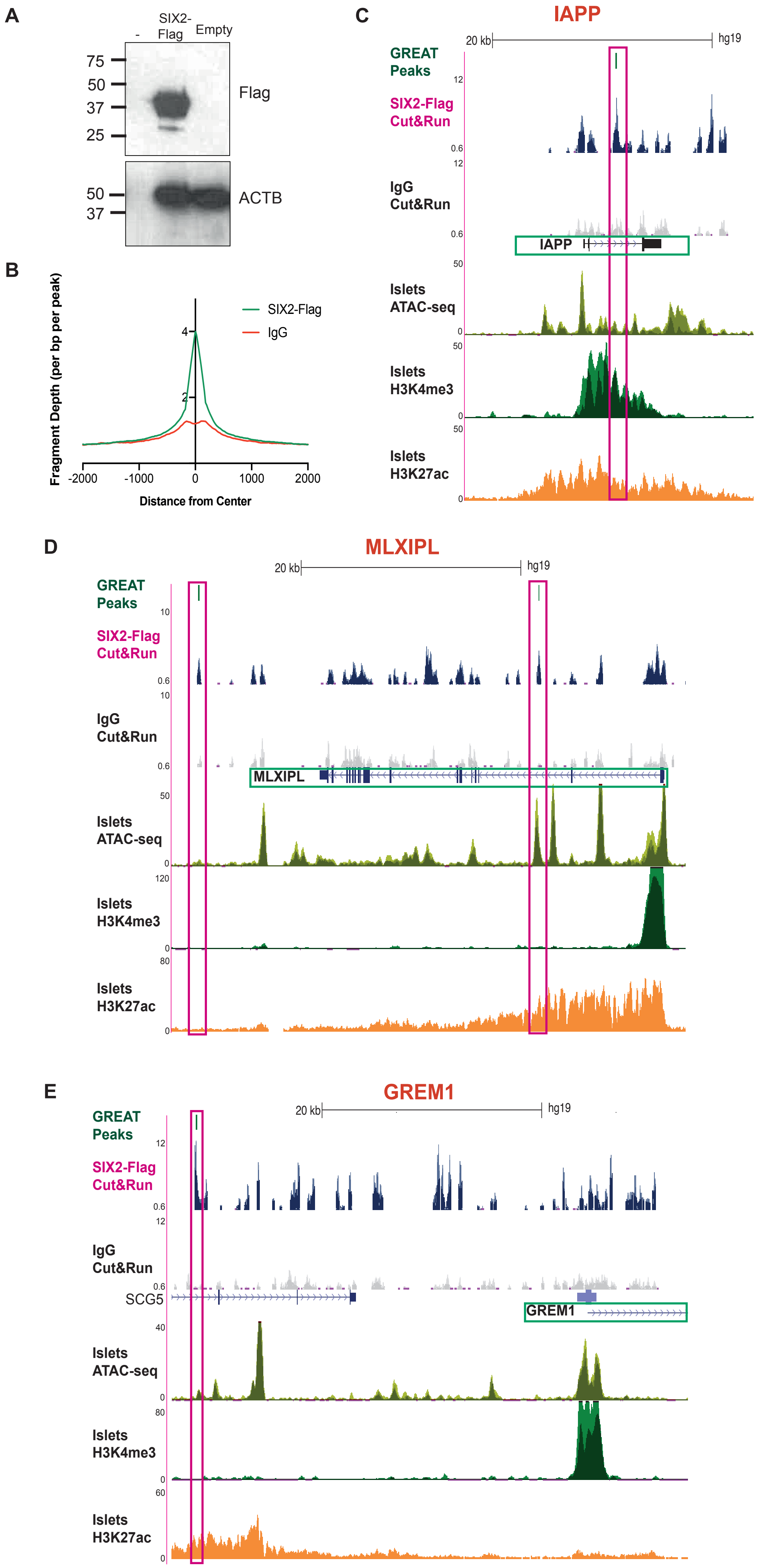

Supplemental Fig. 5

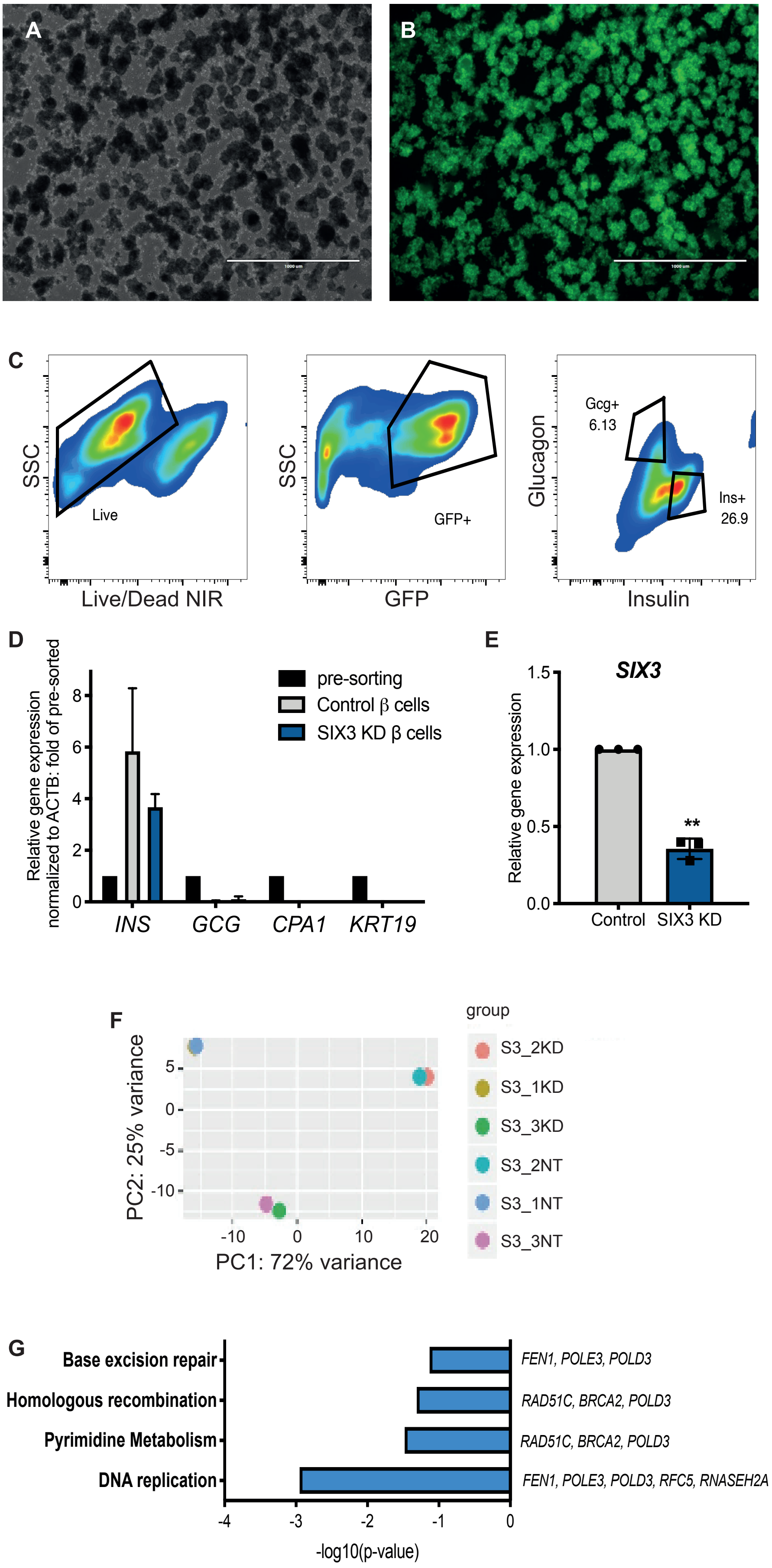

Supplemental Fig. 6

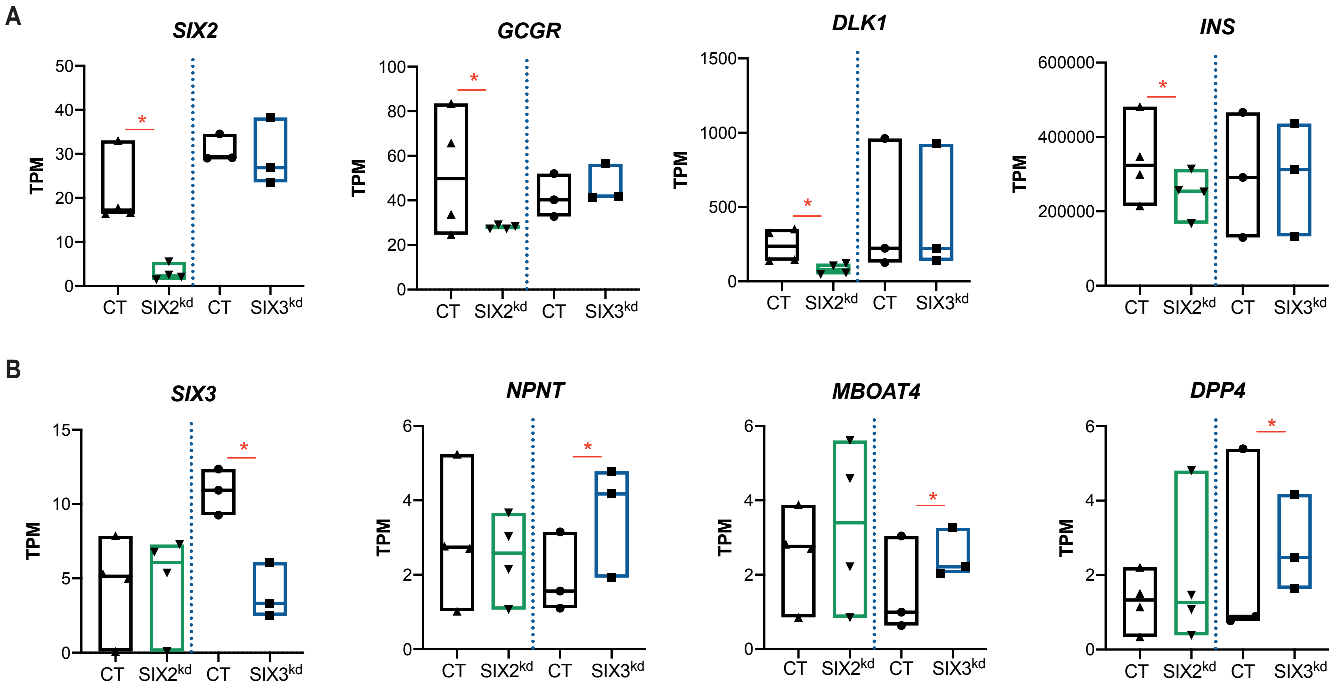

Supplemental Fig. 7

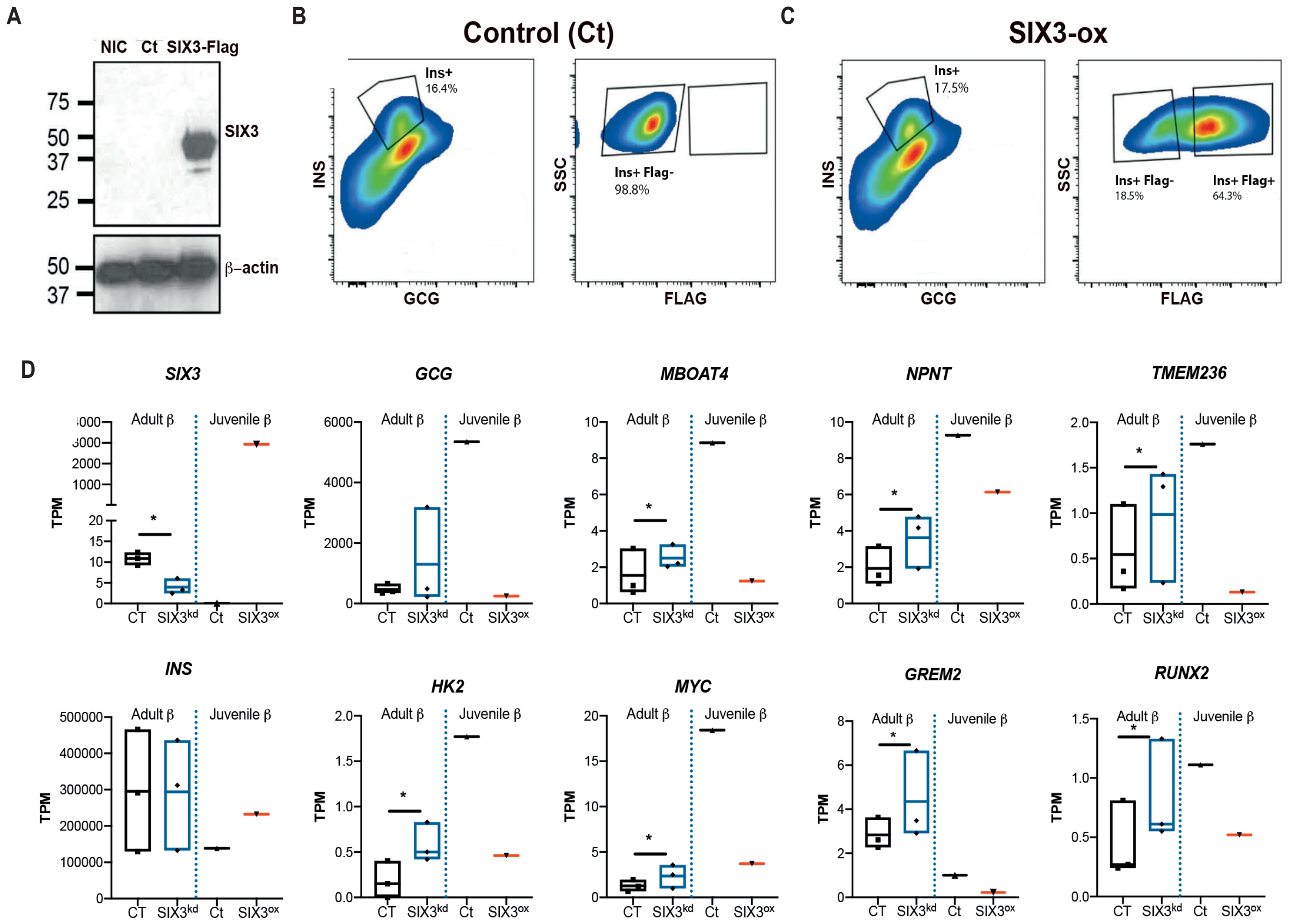

**Table S1.** Information of pancreatic islet donors used in this study.

| Donor prep source | Identification | Sex | Age | BMI | Usage |
|---|---|---|---|---|---|
| IIDP | SAMN08617638 | M | 49 | 34 | SIX3-KD RT-qPCR +GSIS |
| Alberta University | R250 | F | 26 | 25.4 | SIX3-KD RT-qPCR +GSIS |
| IIDP | AEJT193A | | 51 | 29 | SIX3-KD RT-qPCR +GSIS |
| IIDP | AEJE412 | | 55 | 30.1 | SIX3-KD RT-qPCR +GSIS |
| Alberta University | R267 | F | 64 | 23.7 | SIX3-KD RT-qPCR +GSIS |
| Alberta University | R268 | F | 48 | 29.2 | SIX3-KD/SIX2-KD RT-qPCR +GSIS |
| IIDP | SAMN09370567 | M | 32 | 28.5 | SIX3-KD/SIX2-KD RT-qPCR +GSIS |
| IIDP | SAMN09862214 | M | 31 | 31.8 | SIX3-KD/SIX2-KD RT-qPCR +GSIS |
| Alberta University | R286 | M | 41 | 20.4 | SIX3-KD/SIX2-KD RT-qPCR +GSIS |
| AFAO295 | SAMN08611213 | F | 63 | 24.97 | SIX3-KD/SIX2-KD RT-qPCR +GSIS |
| UCSF | rHIP-122 | M | 58 | 27.82 | SIX3-KD/SIX2-KD RT-qPCR +GSIS |
| UCSF | rHIP-144 | F | 47 | 23.51 | SIX3-KD/SIX2-KD GSIS |
| IIDP | SAMN16191825 | | 25 | 31.3 | SIX3-KD/SIX2-KD GSIS |
| IIDP | SAMN15942269 | F | 63 | 26.9 | SIX3-KD/SIX2-KD GSIS<br>Whole islet RT-qPCR -non-diabetic |
| IIDP | SAMN09867385 | M | 24 | 27.2 | SIX3-KD RNA-seq |
| IIDP | SAMN10079665 | M | 25 | 31 | SIX3-KD RNA-seq |
| Alberta University | R288 | F | 69 | 27.7 | SIX3-KD RNA-seq |
| Alberta University | R318 | M | 54 | 20.5 | SIX2-KD RNA-seq |
| UCSF | RHIP-133 | F | 26 | 25 | SIX2-KD RNA-seq |
| Pittsburgh | | M | 24 | 31.8 | SIX2-KD RNA-seq |
| Alberta University | R326 | M | 26 | 27 | SIX2-KD RNA-seq |
| IIDP | SAMN13938639 | F | 50 | 39.2 | RIP-SIX2-Flag CUT&RUN |
| Alberta University | R388 | M | 45 | 33.8 | RIP-SIX2-Flag CUT&RUN |
| IIDP | SAMN16427178 | F | 42 | 31.2 | RIP-SIX2-Flag CUT&RUN |
| CAOP | AHAX150 | F | 3 | n.a | SIX3-overexpression, Beta cell RT-qPCR |
| IIAM | 18 month old | | 18 month | n.a | SIX3-ox, Beta cell RT-qPCR and RNA-seq |
| Alberta University | R282 | M | 57 | 26.4 | Whole islet RT-qPCR-non-diabetic |
| Alberta University | R283 | M | 22 | 22.5 | Whole islet RT-qPCR -non-diabetic |
| Alberta University | R357 | M | 64 | 24.3 | Whole islet RT-qPCR -non-diabetic |
| Alberta University | R348 | F | 43 | 16.4 | Whole islet RT-qPCR -non-diabetic |
| IIDP | SAMN10977276 | M | 52 | 27.2 | Whole islet RT-qPCR -non-diabetic |
| CAOP | AHHQ227 | F | 18 | 34.9 | SIX3-KD/SIX2-KD GSIS; Whole islet RT-qPCR -non-diabetic |
| Alberta University | R347 | M | 57 | 27.9 | Whole islet RT-qPCR -T2D-diabetic |
| NDRI | AGJU173 | F | 53 | | Whole islet RT-qPCR -T2D-diabetic |
| HPAP | AHBD-489 | F | 50 | 30.5 | Whole islet RT-qPCR -T2D-diabetic |
| HPAP | AHGP286 | M | 40 | 37 | Whole islet RT-qPCR -T2D-diabetic |
| HPAP-PANC-DB | HPAP-003 | M | 29 | 24.5 | Non diabetic Beta, alpha cells RNA-seq, islet perifusion |

| | | | | | |
|---|---|---|---|---|---|
| HPAP-PANC-DB | HPAP-006 | M | 46 | 19.1 | Non diabetic Beta, alpha cells RNA-seq, islet perifusion |
| HPAP-PANC-DB | HPAP-008 | F | 24 | 31.9 | Non diabetic Beta, alpha cells RNA-seq, islet perifusion |
| HPAP-PANC-DB | HPAP-004 | F | 24 | 32.2 | Non diabetic Islet perifusion |
| HPAP-PANC-DB | HPAP-014 | F | 43 | 30.9 | Non diabetic Beta, alpha cells RNA-seq |
| HPAP-PANC-DB | HPAP-017 | M | 30 | 23.7 | Non diabetic Beta, alpha cells RNA-seq, islet perifusion |
| HPAP-PANC-DB | HPAP-001 | M | 47 | 32.2 | T2-Diabetic Beta, alpha cells RNA-seq |
| HPAP- PANC-DB | HPAP-007 | F | 65 | 42.6 | T2-Diabetic Islet perifusion |
| HPAP- PANC-DB | HPAP-010 | F | 42 | 36.8 | T2-Diabetic Beta, alpha cells RNA-seq, perifusion |
| HPAP-PANC-DB | HPAP-013 | F | 28 | 41.6 | T2-Diabetic Beta, alpha cells RNA-seq, perifusion |

**Table S7.** *SIX2*$^{kd}$ in primary β-cells results in de-regulation of genes whose expression changes with age in β-cells (see methods for details).

| ***SIX2*$^{kd}$ downregulated genes (*P*<0.05) enriched in adult β-cells (*P*<0.01)** | | | | ***SIX2*$^{kd}$ upregulated genes (*P*<0.05) enriched in juvenile β-cells (*P*<0.01)** | | |
|---|---|---|---|---|---|---|
| AACS | FAM107A | NOMO1 | SORL1 | ACAT2 | PARD6B | POMZP3 |
| ABLIM1 | FAM3B | NPAS2 | SPINT2 | ARHGAP11B | PBX2 | PTGFR |
| ACAT1 | FAM47E | NRCAM | SPTBN4 | ARIH2OS | PCGF6 | ZBTB2 |
| ADAP2 | FBLN7 | OLFM1 | ST14 | ARSK | PHF19 | ZNF581 |
| ADCYAP1 | FBXO2 | OMA1 | STEAP2 | ATG2A | PHLDA1 | |
| ADSSL1 | FBXO44 | P2RX2 | STEAP3 | AURKA | PIM2 | |
| AGT | FRRS1L | P4HA2 | STUB1 | BAK1 | PIP5K1A | |
| AKR7A3 | GALNT11 | PAM | STXBP6 | BAZ1A | PLAGL2 | |
| ALDOA | GCGR | PAPSS2 | SULT4A1 | BTK | POLE4 | |
| ANKH | GDA | PCDHA10 | SUN1 | C12orf4 | PPM1D | |
| ANKRD24 | GHDC | PCDHA6 | SUSD4 | C16orf72 | PRR14 | |
| ANKRD34C | GLT8D2 | PCDHB5 | SYNE2 | C19orf54 | PTGS2 | |
| ANXA7 | GOLGA8A | PCDHGA4 | SYNPO | CACNB1 | PYCR1 | |
| APCDD1L | GOLGA8B | PCDHGA8 | SYT16 | CCL20 | RAB42 | |
| APH1A | GPRASP1 | PCDHGB7 | TBC1D8 | CCL5 | RACGAP1 | |
| AQP3 | GREM1 | PEBP1 | TGOLN2 | CCNA2 | RASGRP3 | |
| ATP6V0E2 | GRIA2 | PFKP | THBD | CCNB2 | RASSF3 | |
| BNIP3 | GRIA3 | PIGZ | TMEM132D | CCNYL1 | RET | |
| C1orf115 | GRIPAP1 | PLCB1 | TMEM176B | CD3EAP | RFX7 | |
| C1orf127 | GRM1 | PLOD2 | TMEM246 | CDC25A | RGMA | |
| C4orf3 | GSN | PLXND1 | TMEM255A | CDCA7L | RMI2 | |
| C7orf50 | HGD | PNMA2 | TMEM59 | CDKN2AIP | SIAH2 | |
| CACNA1C | HHAT | POP4 | TMEM74B | CDR2L | SLC16A4 | |
| CACNA1D | HIBADH | PPIP5K1 | TMOD1 | CENPM | SLC38A2 | |
| CAPN13 | HIVEP3 | PPL | TMX4 | CITED2 | SMC4 | |
| CAPN2 | HOOK2 | PPP2R2C | TNS1 | CLDN15 | SNAI1 | |
| CBX7 | HOPX | PRDX1 | TPD52 | CLSPN | SOD2 | |
| CCDC149 | HPRT1 | PRDX4 | TSPAN13 | CXorf38 | SPNS1 | |
| CCDC85C | HSF4 | PRKAG2 | TSPAN7 | DAPP1 | SRF | |
| CD5 | HSPA12A | PRMT2 | TUBA1A | DCAF12 | SRSF3 | |
| CD9 | IAPP | PROCR | TXNDC5 | DCUN1D3 | STK17B | |
| CD99 | IDS | PRSS23 | TYW3 | ESPL1 | SUV39H2 | |
| CDKN1A | IGFBP3 | PRUNE2 | UCHL1 | EXOSC5 | TCEANC2 | |
| CDKN1C | IGIP | PSAP | UNC79 | FAM189B | TMC6 | |
| CHGB | IGSF21 | PTPN3 | VAT1L | FAM83D | TMEM170A | |
| CHRM3 | IL17RB | PTPRN | VWA5A | FANCA | TMPO | |
| CLCN3 | IL20RA | QPCT | VWDE | FOSL2 | TNFRSF10C | |
| CLU | ITPR3 | RAB17 | WDR45B | GAS2L3 | TONSL | |
| CMTM4 | JAKMIP3 | RAP1GAP2 | WFS1 | H2AFX | TOR4A | |
| CMTM8 | KCND3 | RASGRF1 | WNT4 | HIST1H2BK | TP53 | |
| CNKSR2 | KCNK1 | RIC3 | WSCD2 | HIST1H4H | TPX2 | |
| COG2 | KCNK17 | RIN2 | WWC3 | HIST2H2AA4 | TRIAP1 | |
| CPB1 | KCNK3 | RUFY3 | XDH | HMGCR | TXNIP | |
| CPE | KIRREL3 | RYR2 | XKR4 | IFFO2 | USP42 | |
| CPLX1 | KLHL36 | SCD5 | ZBTB22 | IFITM3 | VNN3 | |
| CPLX2 | LAMA5 | SCG5 | ZFYVE28 | IL1RN | WDR47 | |
| CPNE4 | LGI3 | SCGB2A1 | ZNF395 | ISG20L2 | WDR62 | |
| CRYL1 | LIMCH1 | SCN1B | AK4 | KBTBD2 | XAF1 | |

| | | | | | |
|---|---|---|---|---|---|
| CTAGE6 | LPIN3 | SCN8A | APBA1 | KCNJ2 | ZNF264 |
| CTSB | LRP2BP | SEC11C | CRYZ | KDM2B | ZNF490 |
| DAP | LYRM9 | SEMA5A | DLK1 | KDM6A | ZNF527 |
| DDR1 | MAN1C1 | SETD7 | EFR3B | KIFC1 | ZNF570 |
| DGCR2 | MAPT | SGSM1 | FA2H | KLF16 | ZNF574 |
| DHRS2 | MBP | SH3YL1 | GPX2 | KLF3 | ZNF669 |
| DIRAS3 | MCOLN3 | SHROOM1 | GREM2 | KLF4 | ZNF829 |
| DLGAP4 | MDH1B | SIAE | HIGD1A | KLHL15 | ZNF835 |
| DMKN | ME3 | SIX2 | HIST1H1E | KPNA2 | ZNF850 |
| DOCK9 | METRN | SLC16A12 | KIF26B | LEAP2 | ZSWIM8 |
| DPH6 | MICAL3 | SLC25A27 | KREMEN1 | LSMEM1 | ARL4A |
| DPP7 | MICU2 | SLC35D3 | LPCAT1 | MCL1 | BBC3 |
| DYNLT3 | MLXIPL | SLC35F3 | MALL | MID1IP1 | BRCA2 |
| DZIP1L | MT1E | SLCO5A1 | NFIC | MSMO1 | C1QTNF9 |
| ECHDC2 | MT1F | SMAD9 | SDK2 | MTF1 | C2CD2L |
| EHBP1 | MT1X | SMARCD3 | SLC7A4 | MTG2 | CCDC116 |
| EHF | MUC13 | SMIM5 | SLC9A3R2 | NEB | CCDC168 |
| ENO2 | MYH14 | SMOC1 | TMEM132B | NETO2 | CD300LB |
| ENPP5 | MYO1D | SMPDL3A | TMEM184B | NLRC4 | CDC45 |
| ENTPD3 | MYO5C | SNAP25 | TSKU | NR4A2 | HSPBAP1 |
| ENTPD6 | NBEAL1 | SNCB | TTC7B | NRAS | ITGA2B |
| EPB41L4B | NDRG1 | SNRNP35 | UNC5B | NUSAP1 | LAT |
| EPCAM | NEIL2 | SNTA1 | WSCD1 | OLFML3 | LRRN3 |
| EPHX2 | NFASC | SNX29 | | | |

**Table S8.** Genes showing DE upon *SIX2*$^{kd}$ and *SIX3*$^{kd}$ in the adult β-cell ($P$<0.05)

| | | | | | | | |
|---|---|---|---|---|---|---|---|
| A1CF | DDX55 | ITM2C | RADIL | VWA1 | ZNF829 | SF3A2 | SLIT1 |
| ACCS | EIF5 | KCNA5 | RBM15 | WDR81 | RRM2B | RNF38 | SLC44A2 |
| ACOT7 | FEN1 | KHSRP | RSAD1 | ZIK1 | GLT8D2 | GPRASP1 | TSKU |
| ACSL6 | FFAR4 | KLHL29 | SH2B3 | ZNF547 | GDF15 | CAPN13 | FBLN1 |
| ANKRD40 | FST | LRP1 | SLC9A3R2 | ZNF57 | FXYD6 | PPIP5K1 | PLXDC2 |
| ARFGEF3 | GAREM1 | LRRC75A | SMIM32 | ZNF77 | SYT7 | ZFYVE28 | NEB |
| ARL14 | GDNF | MAST1 | SPEG | ZSWIM4 | RGS11 | ZBTB22 | CCL20 |
| ATP2A3 | GHRL | MAX | SRA1 | TPX2 | SCD5 | SHROOM1 | |
| B4GALNT3 | GPR142 | MPV17L2 | STOM | GAS2L3 | RAP1GAP | NCKAP5 | |
| BRCA2 | GRIN2C | MYO1A | SVOP | CDC37L1 | HILPDA | PFKP | |
| BRINP1 | HGH1 | OLIG1 | TAF5 | ZNF556 | OBSL1 | KCNMB2 | |
| BST2 | HIPK2 | OVGP1 | THOC5 | ZNF296 | SNCB | GREM2 | |
| C4B | HIST2H2AA3 | PCDHA7 | TLE2 | CDR2L | RAP1GAP2 | KCND3 | |
| CACNA1H | HS6ST3 | PHF7 | TMEM132A | PSAT1 | ITPR3 | COL6A1 | |
| CAMK2N1 | IDI1 | PLCH2 | TMEM237 | CD3EAP | TRPM3 | PTPRS | |
| CD300A | IFIT2 | POLE3 | TREX1 | PROCR | NDRG1 | HAP1 | |
| COL27A1 | INSYN1 | PTK7 | TSPOAP1 | CRELD2 | CCDC85C | HGD | |
| CXCL5 | ISG20 | RAD21 | URGCP.MRPS24 | SUPV3L1 | TNS1 | PBX2 | |

**Table S9.** *SIX3*$^{kd}$ in primary $\beta$-cells results in upregulation of genes enriched in juvenile $\beta$-cells (see methods for details).

| ***SIX3***$^{kd}$ **upregulated genes (*P*<0.05) enriched in juvenile $\beta$-cells (*P*<0.01)** | | | | | | |
|---|---|---|---|---|---|---|
| ADD3 | CELF3 | ELF4 | INSR | NFIB | PTPRS | VNN1 |
| ANO6 | CNN2 | EVI2B | IQGAP2 | NPAS4 | SERPING1 | WIZ |
| ANTXR2 | CNOT3 | FAM126A | KCNMB2 | PBX2 | SH3GL1 | ZNF341 |
| ANXA1 | DMD | FBLN1 | KCTD12 | PCGF2 | SLC1A5 | C4B |
| ATG9A | DMTN | FOXP4 | LYZ | PDK2 | SLC44A2 | CD302 |
| BCL6 | DOCK8 | FSTL1 | MVB12B | PHF12 | THBS1 | GHRL |
| BCL7A | DPP4 | HILPDA | MYC | PLXDC2 | TJP2 | INPPL1 |
| CCL20 | DUSP2 | HK2 | NCKAP5 | PLXNA1 | TLE3 | RNF182 |
| CCNG2 | EBF4 | IER5L | NDST2 | PPDPF | TMEM236 | TNRC18 |
| CD248 | EHD2 | INSM1 | NEB | PTPN12 | TSKU | VIT |

**Table S12.** List of genes DE by *SIX2*$^{kd}$ both in adult human β-cells (this study) and SC–β-like cells (Velazco-Cruz et al 2020). DESeq2 was used to calculate DE genes with log2FC (*P*<0.05)

| Genes downregulated post *SIX2*$^{kd}$ in human β-cells and SC–β-like cells | | | | | Genes upregulated post *SIX2*$^{kd}$ in human β-cells and SC–β-like cells | |
|---|---|---|---|---|---|---|
| A1CF | DEPP1 | ISYNA1 | PCDHGA8 | SLC35F3 | AOC3 | PLAGL2 |
| ABCA2 | DEPTOR | ITM2C | PCDHGB7 | SLC4A3 | ARL4A | POLE3 |
| ABCC3 | DHRS2 | JAKMIP3 | PCP4 | SLC4A8 | ARRDC4 | POLR1E |
| ABCC4 | DIRAS3 | KCNA5 | PCSK1N | SLC7A4 | ATP8B2 | POLR3D |
| ABHD14A | DLK1 | KCND3 | PCSK6 | SLC9A3R2 | B3GALNT2 | PRG4 |
| ABHD14B | DMKN | KCNIP2 | PDE9A | SLCO5A1 | BAZ1A | PRMT3 |
| ABLIM1 | DNM1 | KCNJ6 | PFKP | SMARCD3 | BST2 | PROB1 |
| ACADVL | DOCK9 | KCNK1 | PFN2 | SMIM32 | C16orf72 | PRR14 |
| ACCS | DPP6 | KCNK16 | PHLDB2 | SMIM6 | C19orf48 | PRRC1 |
| ACOXL | DSCAML1 | KCNK17 | PIGZ | SMKR1 | C21orf91 | PRSS8 |
| ACSL6 | DZIP1L | KCNMA1 | PIPOX | SMOC1 | CALU | PRX |
| ACTN1 | ECEL1 | KCNMB2 | PLA2G6 | SNAP25 | CCDC134 | PTCH2 |
| ADA2 | EDN3 | KCNQ2 | PLCB1 | SNCB | CCDC17 | PTGFR |
| ADAMTS2 | ELF3 | KIAA1211L | PLCE1 | SNTA1 | CCNB1IP1 | PTGS2 |
| ADCY1 | EMB | KIF12 | PLD3 | SORL1 | CCSAP | PYCR1 |
| ADCYAP1 | ENO2 | KIF5C | PLEKHB1 | SPATA20 | CD274 | RALA |
| ADSSL1 | ENTPD1 | KIRREL3 | PLEKHG6 | SPON2 | CDC25A | RASSF1 |
| AK4 | EPB41L1 | KLHL29 | PLXND1 | SPTBN4 | CDCA7L | RCC1 |
| ALDOA | EPB41L4B | LGI3 | PNMA2 | SRGAP3 | CEBPB | RCL1 |
| ANGPTL4 | EPHX2 | LIMCH1 | PNPLA4 | SSTR2 | CEP85 | RGMA |
| ANKRD24 | F12 | LINGO2 | POMGNT1 | ST8SIA3 | CFAP77 | RIPK2 |
| ANKRD34C | FAM131C | LMO1 | PPL | STEAP2 | CHKA | SAR1A |
| ANKRD37 | FAM228B | LPIN3 | PPP1R36 | STEAP3 | CITED2 | SESN2 |
| ANKRD6 | FAM229B | LPP | PPP2R2C | STXBP1 | CLDND2 | SHMT1 |
| APBA1 | FBXO2 | LRFN2 | PPY | SULF2 | CMYA5 | SLC22A1 |
| APPL2 | FBXO44 | LRP2BP | PRAG1 | SULT4A1 | COA7 | SLC30A1 |
| ARFGEF3 | FFAR1 | MAGI1 | PRKACA | SUSD4 | CREB3 | SLC38A2 |
| ARSD | FFAR4 | MALL | PRKAG2 | SYNGR1 | CYTH3 | SNX8 |
| ASRGL1 | FGF14 | MAN1C1 | PRPH | SYNPO | DCLRE1B | SRF |
| ATP2B4 | FGFR1 | MAP1LC3A | PRPS2 | SYT16 | DDX20 | STX6 |
| ATP6V0E2 | FGGY | MAPK10 | PRSS22 | SYT17 | DDX55 | SUV39H2 |
| B3GNT3 | FRRS1L | MAPRE3 | PSD | SYT7 | DNAH11 | TAF1D |
| BAALC | FSTL4 | MDH1B | PSD4 | TAGLN3 | DUSP14 | TAPT1 |
| BABAM2 | FXYD3 | ME3 | PTGDR2 | TCERG1L | EIF5 | TEAD2 |
| BACE2 | FXYD6 | MEIS3 | PTK2B | TCTN1 | ESAM | TGIF2 |
| BAIAP3 | FYB2 | METRN | PTPRN | TENM1 | ESPL1 | THAP9 |
| BDKRB2 | GABARAPL2 | MGAT4C | PTPRS | THBD | FAM160A1 | THUMPD2 |
| BLOC1S6 | GABRA2 | MISP | PTPRT | TIAM1 | FLCN | TMEM136 |

| | | | | | | |
|---|---|---|---|---|---|---|
| BRINP1 | GALNT10 | MLXIPL | PXK | TM4SF4 | FOSL2 | TMEM167B |
| BTN3A3 | GAMT | MPP2 | QDPR | TMEM125 | FST | TMEM39A |
| C1orf127 | GCG | MRAP2 | QPCT | TMEM132A | HIST1H2BK | TMPO |
| C22orf42 | GCK | MROH7 | RAB26 | TMEM132D | HIST1H4J | TOR4A |
| C9orf16 | GDA | MS4A8 | RAB31 | TMEM158 | HIST3H2A | TP53 |
| CACNA1C | GDPD5 | MSRB2 | RAB3B | TMEM163 | HMOX1 | VPS37B |
| CAMK1D | GJB1 | MTA3 | RAB4B | TMEM255A | HRK | WDR90 |
| CAMK2B | GLT8D2 | MYH10 | RAB9B | TMEM42 | ICK | X9.Mar |
| CAMK2N1 | GNB5 | MYL7 | RABL3 | TMEM61 | IFFO2 | ZBTB24 |
| CAPN2 | GPD1 | MYO1D | RASA4 | TMEM74B | ISG20L2 | ZBTB5 |
| CASR | GPR148 | MYO5B | RASGRF1 | TMOD1 | KBTBD8 | ZHX1 |
| CBX7 | GPRASP1 | MYT1L | RBP4 | TMPRSS6 | KIF24 | ZIK1 |
| CCDC159 | GRIA2 | NBEAL1 | RDX | TMX4 | KIF27 | ZNF267 |
| CCKBR | GRIN3B | NCAM1 | RGS11 | TNS2 | KLF3 | ZNF474 |
| CCND2 | GRM1 | NCKAP5 | RHOU | TP53I11 | KLHL21 | ZNF551 |
| CCNI | GSN | NDRG1 | RIMBP2 | TPM4 | KMT5A | ZNF57 |
| CDH22 | GSTM2 | NECAB2 | RTN1 | TRIM2 | LRRC66 | ZNF581 |
| CDKN1A | HAP1 | NFAM1 | RTN4RL1 | TRMT9B | LYAR | ZNF749 |
| CELF4 | HEPACAM2 | NFASC | RYR2 | TSPAN1 | LYG1 | ZNF813 |
| CERK | HOPX | NLRP1 | S100B | TSPAN7 | MCM2 | ZNF841 |
| CHD3 | HSD3B7 | NPAS2 | SARDH | TSPOAP1 | METTL1 | ZNF850 |
| CHDH | HSF4 | NPDC1 | SCG5 | TTLL6 | MGME1 | |
| CHGB | HSPA12A | NPW | SCN1B | TUB | MMP14 | |
| CLDN4 | HSPD1 | NRBP2 | SDK2 | UNC5A | MMS22L | |
| CLU | IAPP | NRXN2 | SFTPD | UNC5B | MPV17L2 | |
| COL6A1 | IDNK | NSG1 | SHISA4 | VAT1L | MRAS | |
| COL9A2 | IDS | OLFM1 | SHISA6 | VTN | MRE11 | |
| CPE | IGFBP3 | OMA1 | SHROOM2 | VWA5A | MSANTD2 | |
| CPLX1 | IGFBP7 | PACRG | SIGIRR | WNT4 | NETO2 | |
| CPLX2 | IGSF21 | PACSIN1 | SIGLEC15 | WSCD2 | NFKBIB | |
| CREG2 | IL11RA | PALM2 | SIL1 | X1.Mar | NRAS | |
| CST3 | IL17RB | PAX6 | SIX2 | XKR4 | NUS1 | |
| CTSB | IL1R1 | PCBP4 | SLC12A8 | ZDHHC22 | ORC6 | |
| CYBA | IL20RA | PCDHA10 | SLC16A12 | ZFYVE28 | PAIP2B | |
| DACH2 | IMPDH1 | PCDHB10 | SLC18A2 | ZNF860 | PEX5 | |
| DCTN1 | INS | PCDHB13 | SLC22A17 | | | |
| DCX | INSYN1 | PCDHB14 | SLC25A27 | | | |
| DDC | IRX3 | PCDHB6 | SLC35D3 | | | |

### Supplemental Figure Legends

**Figure S1. Pseudoislets exhibit re-aggregation of the principal islet cell-types after lentiviral transduction**
(A-L) Immunostaining of INS (red) and GCG (white) paired with GFP (green) in human pseudoislets five days after transduction with (A-D) Control, (E-H) *SIX2*$^{kd}$ and (I-L) *SIX3*$^{kd}$ lentiviruses. (M-X) Immunostaining of INS (white) and STT (red) paired with GFP (green) in human pseudoislets five days after transduction with (M-P) Control, (Q-T) *SIX2*$^{kd}$ and (U-X) *SIX3*$^{kd}$ lentiviruses. Scale bars, 20 µm.

**Figure S2. β-cells *SIX2*$^{kd}$ RNAseq**
(A) *SIX2* mRNA expression in sorted β-cells of control (grey bar) and *SIX2*$^{kd}$ (green bar) (n=4 independent donors). (B) Insulin, Glucagon CPA-1 and KRT19 mRNA expression in pre-sorted (black bars), GFP+ Insulin+ control (grey bars) and *SIX2*$^{kd}$ (green bars) pseudoislet cells, data normalized to human beta actin. (C) Detailed FACS scheme used to sort GFP+ β−cells of Control and *SIX2*$^{kd}$ pseudoislets: single, live, GFP$^{+}$ cells were used for intracellular staining with INS and GCG antibodies, and INS$^{+}$ cells were used for RNA extraction and RNA-Seq library building: see methods. (D) PCA plot showing variance due to knockdown and donor conditions (n=4 independent donors). (E) Heatmap of the sample-to-sample distances of all RNA-Seq samples used in this experiment. The scale bar indicates the euclidean distance. The data is presented as mean, error bars represent the standard error. Two-tailed t tests were used to generate *P* values. *** $P<0.0001$

**Figure S3. *SIX2*$^{kd}$ in β-cells results in a distinct set of differentially expressed genes compared to SC β-like cells**: (A) *SIX2*$^{kd}$ downregulated genes (B) *SIX2*$^{kd}$ upregulated genes. For both datasets, DE genes were calculated using DEseq2, $P<0.05$ (see methods).

**Figure S4. SIX2-Flag CUT&RUN in adult human islets.** (A) Western blot using anti-Flag antibody on protein lysates from either SIX2-Flag or control (Empty construct) human islets. (B) Histogram plotting averaged reads showing enrichment of read densities in the peak centers for the SIX2-Flag libraries. (C-E) Tracks showing SIX2-Flag genomic regions associated to (D) IAPP, (E) MLXIPL, (F) GREM1. Accessible chromatin regions in human islets are shown by ATAC-seq, H3K27ac and H3K4me3 ChiP-seq tracks of whole human islets. SIX2 CUT&RUN (SIX2 C&R) peaks are shown in pink boxes, and regulated genes highlighted in green boxes.

**Figure S5. β-cells *SIX3*$^{kd}$ RNAseq**: (A,B) Human pseudoislets five days after transduction with *SIX3*$^{kd}$ lentiviruses: (A) bright field; (B) blue light (488 nm), scale bars = 1000 µm. (C) Detailed FACS scheme used to sort *SIX3*$^{kd}$ GFP$^{+}$ β−cells: live, GFP$^{+}$, INS$^{+}$ cells were sorted for RNA extraction and RNA-Seq library building. (D) *SIX3* mRNA expression in sorted β-cells of control (grey bar) and *SIX3*$^{kd}$ (blue bar) (n=3). (E) Insulin, Glucagon CPA-1 and KRT19 mRNA expression in pre-sorted (black bars), GFP+ Insulin+ control (grey bars) and *SIX3*$^{kd}$ (blue bars) pseudoislet cells (n=3 independent donors), data normalized to human beta actin. (F) PCA plot showing variance due to knockdown and donor conditions. (n=3). (G) KEGG pathways enriched in genes downregulated in β-cells post-*SIX3*$^{kd}$. The data is presented as mean, error bars represent the standard error. Two-tailed t tests were used to generate *P* values. ** $P<0.05$

**Figure S6. SIX2 and SIX3 targets are distinct:** Boxplots displaying normalized TPM counts of (A) selected S*IX2*$^{kd}$ targets in the adult β-cell; the expression of which is not affected by *SIX3*$^{kd}$, control (grey bars), S*IX2*$^{kd}$ (green bars), S*IX3*$^{kd}$ (blue bars). (B) selected S*IX3*$^{kd}$ targets in the adult β-cell; the expression of which is not affected by *SIX2*$^{kd}$, control (grey bars), S*IX2*$^{kd}$ (green bars), S*IX3*$^{kd}$ (blue bars). Box plots show the mean. *, *P*<0.05

**Figure S7. *SIX3* overexpression in juvenile β-cells**: (A) Western blot using anti-SIX3 antibody on protein lysates from either non infected control (NIC), infected Control (Ct, Empty construct) or SIX3-Flag overexpressed human islets. (B-C) FACS scheme used to sort Flag+ SIX3-ox juvenile β–cells, using intracellular staining with INS and GCG antibodies and intranuclear staining with Flag antibody, for the Empty Control (Ct) and *SIX3*-Flag (SIX3-ox) overexpressed human islets. (D) Boxplots displaying normalized TPM counts of *SIX3*, *INS* and selected S*IX3* targets in the adult β-cell; the expression of which is inversely affected by SIX3-ox in juvenile β-cells (n=1), control (black bars), S*IX3*$^{kd}$ (blue bars), S*IX3*$^{Ox}$ (red bars). Box plots show the mean. *, *P*<0.05
